# Expert-guided protein Language Models enable accurate and blazingly fast fitness prediction

**DOI:** 10.1101/2024.04.24.590982

**Authors:** Céline Marquet, Julius Schlensok, Marina Abakarova, Burkhard Rost, Elodie Laine

**Author notes:** Céline Marquet and Julius Schlensok contributed equally to this work.

## Abstract

Exhaustive experimental annotation of the effect of all known protein variants remains daunting and expensive, stressing the need for scalable effect predictions. We introduce VespaG, a blazingly fast missense amino acid variant effect predictor, leveraging protein Language Model (pLM) embeddings as input to a minimal deep learning model. To overcome the sparsity of experimental training data, we created a dataset of 39 million single amino acid variants from the human proteome applying the multiple sequence alignment-based effect predictor GEMME as a pseudo standard-of-truth. This setup increases interpretability compared to the baseline pLM and is easily retrainable with novel or updated pLMs. Assessed against the ProteinGym benchmark (217 multiplex assays of variant effect - MAVE - with 2.5 million variants), VespaG achieved a mean Spearman correlation of 0.48±0.02, matching top-performing methods evaluated on the same data. VespaG has the advantage of being orders of magnitude faster, predicting all mutational landscapes of all proteins in proteomes such as *Homo sapiens* or *Drosophila melanogaster* in under 30 minutes on a consumer laptop (12-core CPU, 16 GB RAM).

**Availability:** VespaG is available freely at https://github.com/jschlensok/vespag. The associated training data and predictions are available at https://doi.org/10.5281/zenodo.11085958.

## Introduction

Proteins are the essential building blocks of life, fulfilling a wide range of vital roles within cells and organisms. Hence, understanding the effect of variations such as point mutations on protein stability and function is crucial for comprehending disease mechanisms (Murray, Laurieri, and Delgoda 2017) and modulating their activities through engineering. Multiplexed assays of variant effect (MAVEs), in particular deep mutational scans (DMS) (Fowler and Fields 2014), have enabled the quantification of mutational outcomes on a much larger scale than ever before. They allow for an in-depth characterization of protein mutational landscapes by assessing the impact of virtually all possible single amino acid substitutions. Nevertheless, conducting experimental assays for entire proteomes remains elusive (Atlas of Variant Effects Alliance n.d.).

Leveraging the power of computational models can help to gain insights into the functional consequences of protein variants and to prioritize them for further experimental validation. However, the sparseness of annotations challenges the development of such models. While many supervised machine learning (ML) methods have proven accurate (Adzhubei, Jordan, and Sunyaev 2013; Gray et al. 2018; Hecht, Bromberg, and Rost 2015), they are inherently biased towards the limited number of proteins characterized by MAVEs or having annotated disease-associated variants (Atlas of Variant Effects Alliance n.d.; Livesey and Marsh 2023). As a result, different methods tend to correlate highly for the tiny subset of experimental data, while their predictions for, e.g., all possible mutations in the human proteome correlate very poorly (Hecht, Bromberg, and Rost 2013; Mahlich et al. 2017). Prediction methods are also sensitive to the noise and uncertainty in these data. MAVE annotations, for instance, may vary substantially across experiments, even when measuring the same phenotype for the same protein (Reeb, Wirth, and Rost 2020). These difficulties have stimulated a growing interest in unsupervised or weakly supervised methods predicting variant effects by only exploiting information from protein sequences observed in nature (Ng and Henikoff 2003).

Among the best-performing unsupervised methods, GEMME explicitly models the evolutionary history of protein sequences (Laine, Karami, and Carbone 2019; Notin et al. 2023). Starting from a multiple sequence alignment (MSA), it determines how protein sites are segregated along the topology of phylogenetic trees to quantify the sensitivity of each site to mutations and the number of changes required to accommodate a substitution. It relies on only a few biologically meaningful parameters and is robust to low variability in the input MSA. GEMME proved instrumental for investigating the interplay between protein stability and function, and elucidating disease mechanisms (Abildgaard et al. 2023; Cagiada et al. 2023; Gersing et al. 2023; Tiemann et al. 2023; Tsuboyama et al. 2023). Combining GEMME with a fast MSA generation algorithm allows for producing proteome-wide substitution score matrices within a few days (Abakarova et al. 2023).

Other methods rely on protein Language Models (pLMs) pre-trained over large databases of raw sequences (Elnaggar et al. 2021; Lin et al. 2023). The log-odds ratios computed from the masked marginal probabilities can already provide highly accurate estimates of mutational effects (Meier et al. 2021)(Atlas of Variant Effects Alliance n.d.; Livesey and Marsh 2023)(Meier et al. 2021). Nevertheless, the quality of the protein sequence representations learned by foundation pLMs is highly variable, and especially poor for viral proteins (Ding and Steinhardt 2024; Elnaggar et al. 2021; Lin et al. 2023; The UniProt Consortium et al. 2023); Notin et al. 2023). While pLM performance can be further boosted through incorporating information about evolutionary conservation, population genetic polymorphism and 3D structures (Su et al. 2024), Cheng et al. 2023; Marquet et al. 2022; Meier et al. 2021; Nijkamp et al. 2022; Notin, Dias, et al. 2022; Truong Jr and Bepler 2023), the computational cost of zero-shot inference over full-length proteins remains high.

Here, we optimized prediction speed by circumventing the computationally expensive masked token reconstruction task and directly mapping pLM embeddings to complete mutational landscapes using the evolutionary-informed model GEMME as a teacher (Fig. 1). To this end, we trained a comparatively shallow (660k free parameters) neural network on top of a pre-trained pLM without computing log-odds ratios to learn GEMME predictions. Our strategy overcomes the bottleneck of sparsely annotated experimental training data. Moreover, it avoids the noise and inconsistencies of the experimental assays. We implemented our approach as a lean tool for fast **V**ariant **E**ffect **S**core **P**rediction without **A**lignments enabled by **G**EMME (<u>VespaG</u>). We assessed prediction performance against over 3 million (M) missense variants across diverse protein families. VespaG performed on par with SOTA methods and in some cases, the student even surpassed the teacher GEMME. As we circumvent the need to compute log-odds ratios of substitution probabilities, VespaG enables proteome-wide predictions in less than a half hour on a standard consumer laptop. We also demonstrated VespaG to generalize across organisms and protein families.

**Figure 1:**
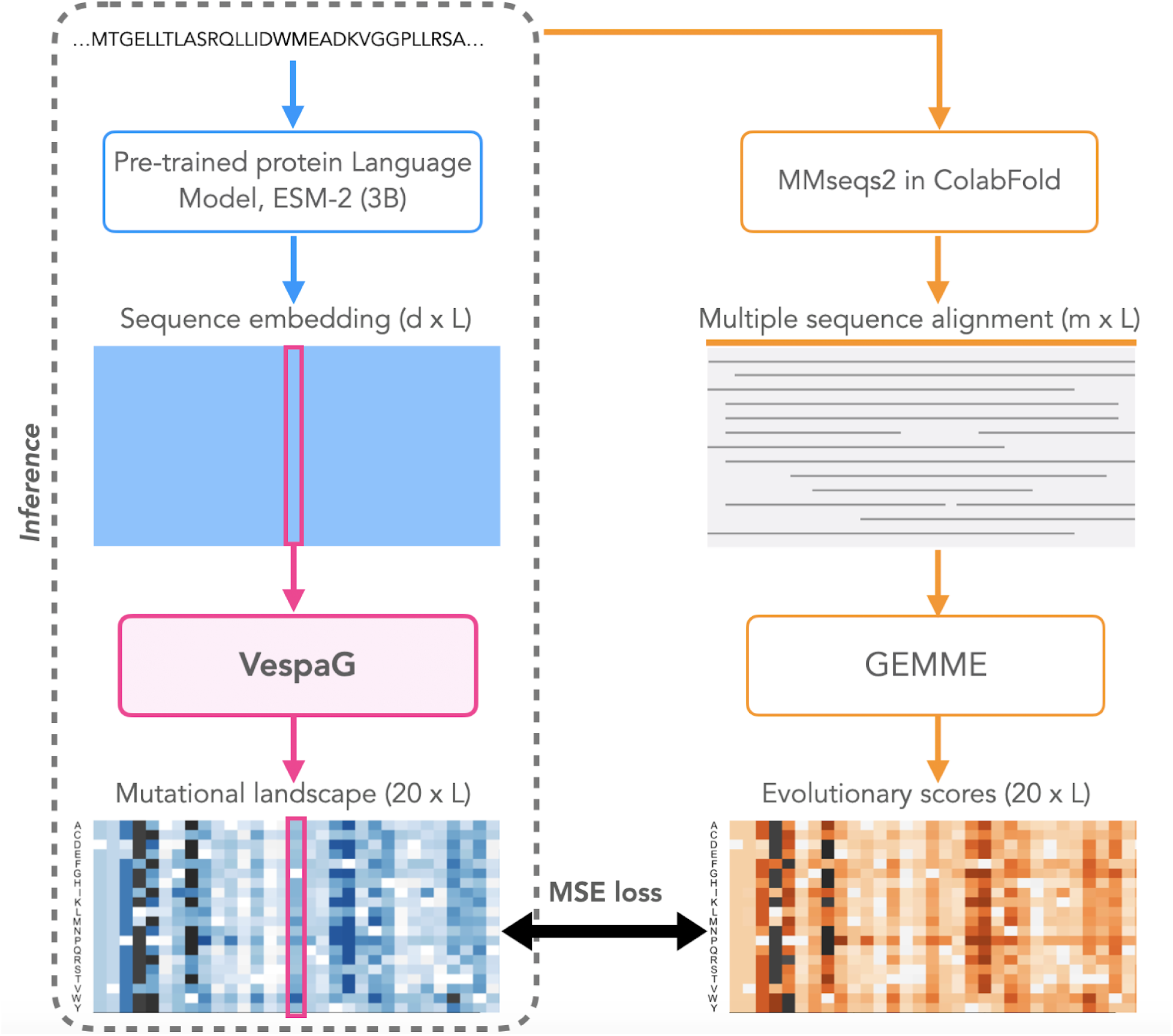
Outline for VespaG’s expert-guided approach. VespaG takes as sole input a d=2560-dimensional vector representation of a wild-type residue in a protein computed by the pre-trained protein language model (pLM) ESM-2 with 3 billion parameters (Lin et al. 2023), and outputs a 20-dimensional vector of predicted mutational outcome estimates. The training loss measures the mean squared error between the predicted estimates and the evolutionary scores computed by GEMME (Laine, Karami, and Carbone 2019). We generate millions of training samples through the MMseqs2-based ColabFold protocol for searching and aligning sequences (Abakarova et al. 2023; Mirdita et al. 2022; Steinegger and Söding 2017). We do not use alignments at inference time (dotted rectangle). VespaG’s framework can be adapted to any pre-trained pLM.

## Results

The method introduced in this work, VespaG, is a feed-forward neural network (FNN) with one hidden layer with 256 hidden units solely inputting sequence embeddings from the protein language model (pLM) ESM-2 (Lin et al. 2023). We trained VespaG on a set of about 5,000 human proteins to learn a mapping between the input pLM embeddings and the evolutionary scores computed by GEMME. The latter served as surrogates for mutational phenotypic outcomes. The proteins used for training represented a non-redundant subset of the human proteome (Methods and Table 1).

**Table 1:** Size statistics of training datasets used. The dataset used for the development of VespaG, dubbed Hum5k, is a redundancy reduced version of the human proteome. To investigate generalizability across organisms, we further analyzed the performance of VespaG trained on several other datasets. These included the redundancy reduced proteomes of *Drosophila melanogaster* (Droso4k) and *Escherichia coli* (Ecoli2k), a redundancy reduced set of all viral proteins in Swiss-Prot (Virus1k), and a redundancy reduced combination of all (All9k).

| Name | Source organism(s) | Number of sequences |  |  |  | Number of residues |  |
| --- | --- | --- | --- | --- | --- | --- | --- |
|  |  | Reference proteome | After Length restriction (Min.25, Max.1024) | After Redundancy Reduction | Confident GEMME predictions | Total | Min., Max., Median protein length |
| **Hum5k** | *Homo sapiens* | 20,357 | 18,043 | 5,886 | 5,305 | 2,010,031 | 36 / 1024 / 328 |
| **Droso4k** | *Drosophila melanogaster* | 13,811 | 12,305 | 4,809 | 4,081 | 1,610,079 | 40 / 1024 / 346 |
| **Ecoli2k** | *Escherichia coli* | 4,403 | 4,284 | 2,544 | 2,333 | 652,257 | 36 / 990 / 243 |
| **Virus1k** | All viral in Swiss-Prot | 17,320 | 15,954 | 4,498 | 1,400 | 401,204 | 29 / 1017 / 228 |
| **All9k** | All of above | 55,891 | 50,586 | 15,175 | 9,616 | 3,291,539 | 29 / 1024 / 286 |

### Influence of hyperparameters and input pLM

We initially considered five different architectures for learning from GEMME (“MSE loss” in Fig. 1), including linear regression and convolutional neural networks, and two foundation pLMs, namely ESM-2 (Lin et al. 2023) and ProtT5 (Elnaggar et al., 2021) (Supporting Online Material, SOM Table S1). Performance was similar for all evaluated models (SOM Fig. S1-2), indicating that both pLMs provide robust results under supervision of GEMME regardless of downstream architecture. As we obtained the best performance with a one-hidden-layer FNN and ESM-2 embeddings against the validation set (random 80/20 split, SOM Fig. S2), we report test set results for this configuration in the following.

### VespaG competitive with state-of-the-art (SOTA)

VespaG predicted mutational outcomes with an average overall Spearman correlation coefficient (ρ) of 0.480±0.021 (±1.96 standard errors, *i.e.*, 95% Confidence Interval, CI; Methods) against all 217 experimental DMS assays from the ProteinGym substitution benchmark (Notin et al. 2023), June 2024). It performed *on par* with the top methods from the ProteinGym leaderboard, including its teacher GEMME, and substantially better than the zero-shot ESM-2 baseline (Fig. 2, SOM Tables S2, S3).

**Figure 2:**
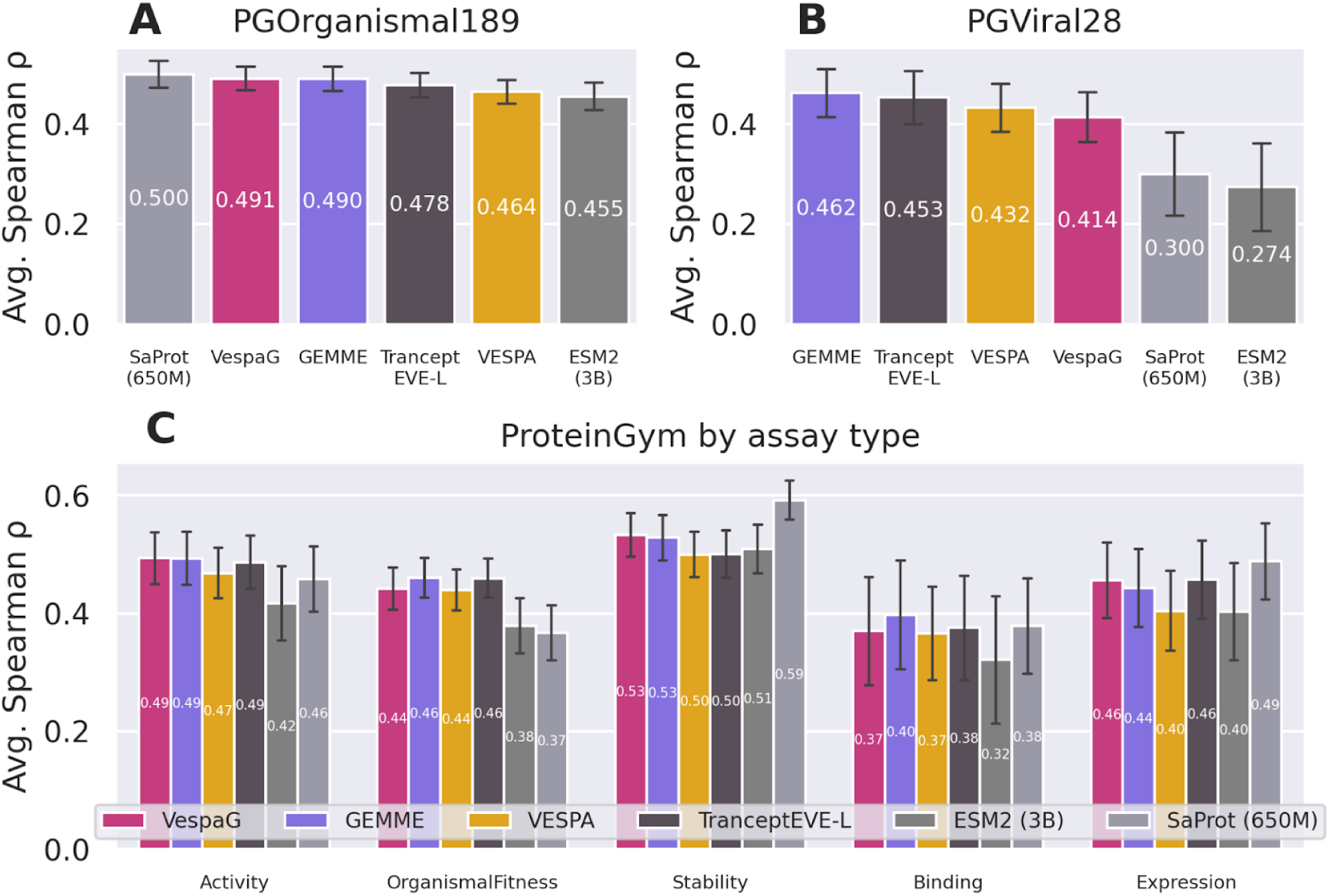
VespaG accuracy on-par with SOTA. Each panel corresponds to a different test set (with partially overlapping proteins between the test sets in C w.r.t A and B) from the ProteinGym substitution benchmark (Notin et al. 2023): (A) *PGOrganismal189*, containing 189 experimental assays for 161 eukaryotic and prokaryotic proteins; (B) *PGViral28*, containing 28 assays for 26 viral proteins; (C) 217 assays in ProteinGym divided into subsets named according to assessed phenotype and number of experiments: *PGActivity43, PGBinding13, PGExpression18, PGFitness77, PGStability66*. For A and B, methods are ordered from best (left) to worst (right), for C we follow a set order. We did not recompute results for TranceptEVE L (Notin, Niekerk, et al. 2022), ESM-2 (Lin et al. 2023), and SaProt (Su et al. 2024), therefore all depicted in shades of gray, and directly extracted the predictions from ProteinGym. The error bars show the 95% confidence interval.

Most of the assays (189 out of 217), were from proteins from eukaryotic and prokaryotic organisms (SOM Fig. S3). On these proteins, VespaG reached an average ρ=0.491±0.024 (Fig. 2A and SOM Table S2, *PGOrganismal189*). Its prediction accuracy exceeded all of the following: zero-shot ESM-2 log-odds ratios between the mutant and wild-type amino acids (Δρ=0.036, one-tailed paired t-test p-value<10^-5^), pLM-based VESPA (Δρ=0.027, p-val<10^-8^) and the ensemble sequence- and MSA-based predictor TranceptEVE L (Δρ=0.013, p-val=0.006). VespaG performed *on par* with its *teacher* GEMME (|Δρ|<0.01, p-val>0.1) and the top-ranked method in ProteinGym (as of June 2024), namely the sequence- and structure-based pLM SaProt (Su et al. 2024). Accuracy varied substantially across different experimental DMS assays (SOM Fig. S3). Yet, VespaG was stable, in the sense that the distribution of its Δρ values with respect to the mean ρ over the six highlighted methods was very narrow and centered around zero (SOM Fig. S5). By contrast, the distributions for the ESM-2 baseline and SaProt were much wider, displaying performance worse than the mean by Δρ<-0.35 for some assays (SOM Fig. S5-S6). Simply put: VespaG appeared to be the most *average* method with the lowest spread between assays (SOM Fig. S6). On a subset of *PGOrganismal189*, dubbed *PGOrganismal66*, we could extend the comparison to the SOTA methods AlphaMissense and PoET (predictions not readily available for other data sets). On this subset, VespaG’s predictive performance, with an average ρ=0.484±0.044, outperformed the pLMs SaProt and ESM-2 (Δρ>0.27, p-val < 0.007), and was slightly better than GEMME and TranceptEVE-L (Δρ>0.07, p-val < 0.04); it was *on par* with PoET (|Δρ|<0.01, p-val>0.1) and comparable to AlphaMissense (Δρ=-0.021, p-val∼10^-3^; SOM Fig. S7).

The experimental DMS assays represent an unbalanced panel of different phenotypes (SOM Fig. S3), namely organismal fitness (*PGFitness77,* highest number of DMS), stability (*PGStability66),* activity (*PGActivity43*), expression (*PGExpression18*), and binding (*PGBinding13)*. Balancing the calculation of the average performance according to the phenotypes’ cardinalities yielded ρ=0.459±0.049 for VespaG, outperforming TranceptEVE L, VESPA, SaProt and ESM-2, and bested only by teacher GEMME (SOM Table S2). VespaG consistently outperformed VESPA and ESM-2. Its relative performance w.r.t. the other methods was reasonably stable across all phenotypes (Fig. 2C, SOM Table S2). In contrast, SaProt performed much better than the other methods on stability (Δρ>0.059), likely due to the fact that it was trained on both sequences and 3D structures, but much worse than all methods except ESM-2 on organismal fitness (Δρ<-0.066). Overall, predictions for stability were the most accurate across all methods followed by activity, organismal fitness, expression, and finally, binding. We observed a large amplitude between the worst average performance, obtained by ESM-2 on binding (ρ=0.312), and the best one, obtained by SaProt on stability (ρ=0.592).

In addition, assessing VespaG performance in function of mutation depth revealed higher Spearman correlations on single missense variants compared to multiple ones (SOM Table S3). We observed a similar trend for all tested predictors, with GEMME consistently yielding the best Spearman correlations. Compared to TranceptEVE L, VESPA, and ESM-2, VespaG’s multi-mutant performance was more stable and more closely aligned to its teacher GEMME, especially for mutations of three or more residues.

### VespaG integrating complementary strengths

We specifically investigated how VespaG improved over its teacher GEMME and its baseline ESM-2 for exploiting the protein sequence universe, dealing with viral proteins, and handling *de novo* proteins.

Only a small subset of 28 DMS assays from ProteinGym concern viral proteins, including eight from *Influenza A virus*, six from *Human immunodeficiency virus*, four from *bacteriophages*, and two from *SARS-Cov-2* (SOM Fig. S3). While VespaG did not match the performance of the top method on this subset, (GEMME, Δρ=-0.048, p-val<10^-4^), it improved substantially over the ESM-2 baseline (Δρ=0.140, p-val<10^-4^) and the sequence- and structure-based pLM SaProt (Δρ=0.113, p-val<10^-3^, Fig. 2B, *PGViral28*; SOM Fig. S5). Thus, VespaG was more accurate on viral proteins than other ESM and SaProt versions (SOM Fig. S8-9). This analysis suggests that supervision via GEMME partially counterbalances the poor quality of pLM embeddings for viral proteins. We obtained similar results on a subset of 21 DMS from ProteinGym’s first iteration (SOM Fig. S7).

The proverbial student VespaG bested the teacher GEMME by a large margin (Δρ > 0.1) for the human protein LYAM1, the murine MAFG, the bacterial proteins DN7A, F7YBW7, ISDH, NUSA, and SBI, the plant RCD1, and the yeast ubiquitin RL40A (SOM Fig. S5). In particular, VespaG correctly identified the glycines G75 and G76 in the top five ubiquitin residues most sensitive to mutations, whereas GEMME incorrectly predicted them as mildly sensitive (Fig. 3). These two residues play essential roles for E1 activation (Mavor et al. 2016). Reciprocally, VespaG agreed with the experiment on the mild tolerance of K27, whereas GEMME predicted this residue as highly sensitive (Fig. 3). We can interpret these discrepancies in light of previous works showing that ubiquitin stands out from the general trends between evolutionary sequence conservation and the experimentally measured tolerance to substitutions (Mavor et al. 2016, 2018; Roscoe et al. 2013). It challenges the common view that high selection pressure implies high mutational sensitivity. Hence, applying this principle on an input MSA, as GEMME does, leads to a limited accuracy (ρ in the 0.36-0.44 range). VespaG’s representation learning-based approach allows overcoming this limitation and capturing key aspects of the peculiar sequence-phenotype ubiquitin relationship (ρ in the 0.48-0.54 range).

**Figure 3:**
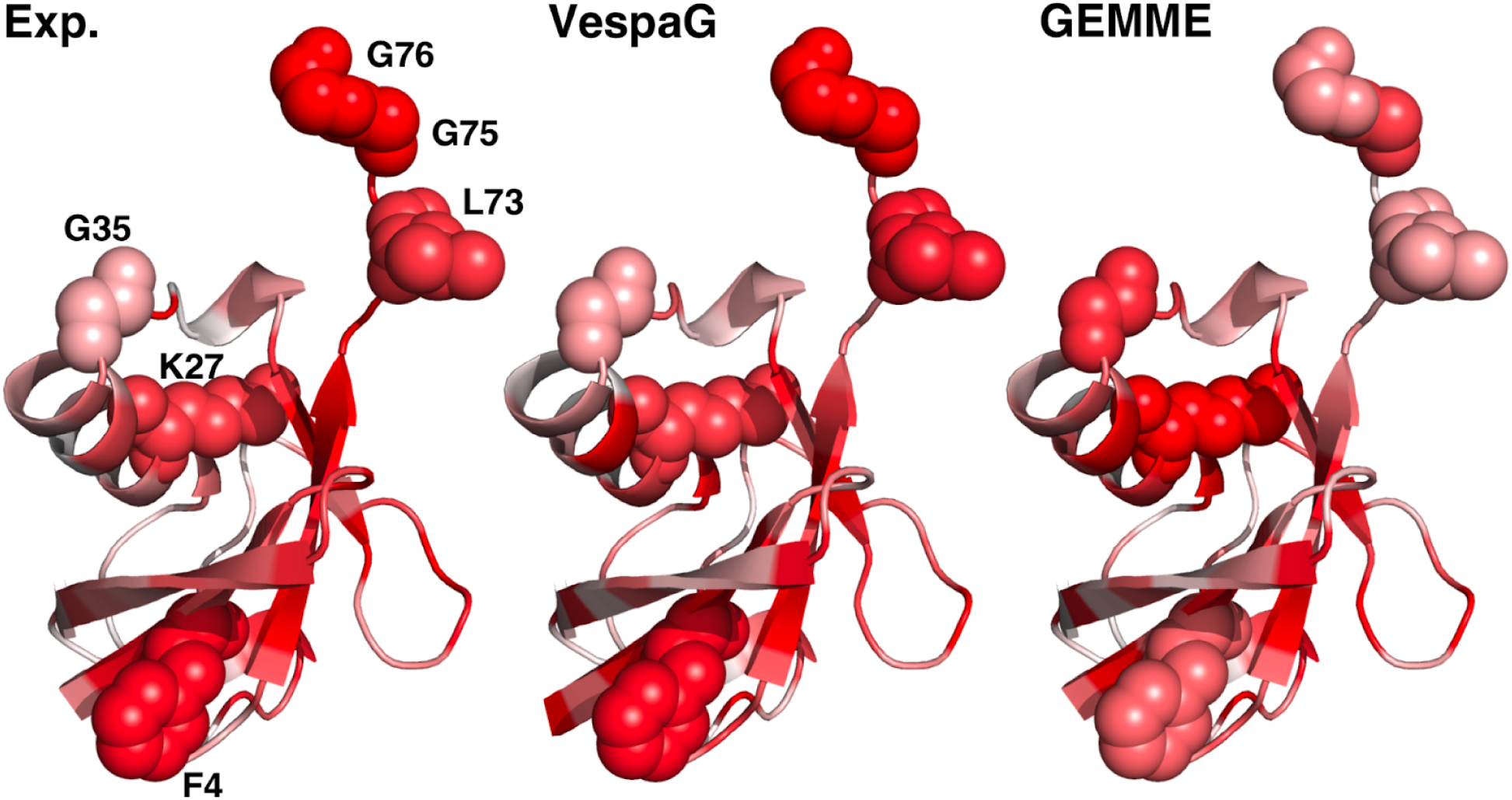
Details of *student* VespaG vs *teacher* GEMME. For the yeast ubiquitin (RL401A_YEAST), we compared experimental measurements (left panel, labeled Exp.; Mavor et al. 2016) with predictions by mapping the per-residue mutational sensitivities onto the 3D structures predicted by AlphaFold2 (AF-P0CH08-F1-model_v4, residues 2 to 76, Jumper et al. 2021). We estimated the extent to which a residue is sensitive to mutations as the rank of its average predicted or measured effect over the 19 possible substitutions. The more reddish the more sensitive. We highlighted six residues (labeled by one-letter amino acid code followed by position in the sequence, *e.g.*, G76: glycine at position 76) for which VespaG agreed with the experiment (rank difference <5) while GEMME strongly disagreed (rank difference >15). The experimental values reflect ubiquitin fitness landscape under normal growth conditions (Mavor et al. 2016).

More generally, VespaG has the advantage of being independent of any alignment, whereas GEMME results may substantially differ depending on the chosen MSA generation protocol. Namely, GEMME Spearman correlations displayed large variations (in the [0.1-0.3] range) for 16 assays when retrieving the input MSAs using ColabFold’s MMseqs2-based strategy (Mirdita et al. 2022) versus taking the ProteinGym MSAs (SOM Fig. S10). The latter were generated with the more sensitive profile Hidden Markov Model search algorithm JackHMMER (Johnson, Eddy, and Portugaly 2010). For almost all these assays (13/16), VespaG achieved a ρ value similar to or higher than the maximum ρ over the two GEMME runs, regardless of the associated MSA generation protocol (SOM Fig. S10). This result suggests that the VespaG framework is at least equivalent to a high-quality MSA-based setup.

Furthermore, VespaG’s independence from alignments makes it applicable to *de novo* proteins. It reached an average Spearman correlation of ρ_nov_=0.404±0.011 on an additional test set of 146 assays reporting mutation-induced thermodynamic folding stability changes for *de novo* designed 40-72 amino acid long protein domains (Tsuboyama et al. 2023) (SOM Fig. S11). By contrast, the baseline zero-shot ESM-2, although technically able to handle *de novo* sequences, yielded an extremely poor average Spearman correlation of 0.034. The teacher GEMME only produced predictions for four *de novo* proteins due to a lack of sufficient input alignments. Its Spearman correlation on this small subset was nearly zero (0.085), compared to 0.393 for VespaG.

### VespaG generalizing across multiple organisms

We further assessed the impact of the training set on VespaG’s predictive performance and ability to transfer knowledge across organisms. Specifically, we retrained from scratch the same architecture with the same hyperparameters on non-redundant sets of ∼4,000 proteins from the insect *Drosophila melanogaster*, ∼2,000 proteins from the bacterium *Escherichia coli*, ∼1,500 proteins coming from several viruses, and ∼9,000 proteins from a combination of all. Training VespaG on a few thousand diverse proteins from a single organism sufficed to generalize across diverse taxa (Fig. 4). Overall, the performance differences for respective taxa were small across organismal training sets (Δρ≤0.1). However, we consistently observed a lower agreement between VespaG predictions and the experiments for viral proteins, compared to other taxa, across all training sets. In particular, exclusively learning from ∼1,500 viral proteins did not improve performance for viral proteins (Fig. 4). Inputting embeddings from other pLMs did not alter this trend (SOM Fig. S2).

**Figure 4:**
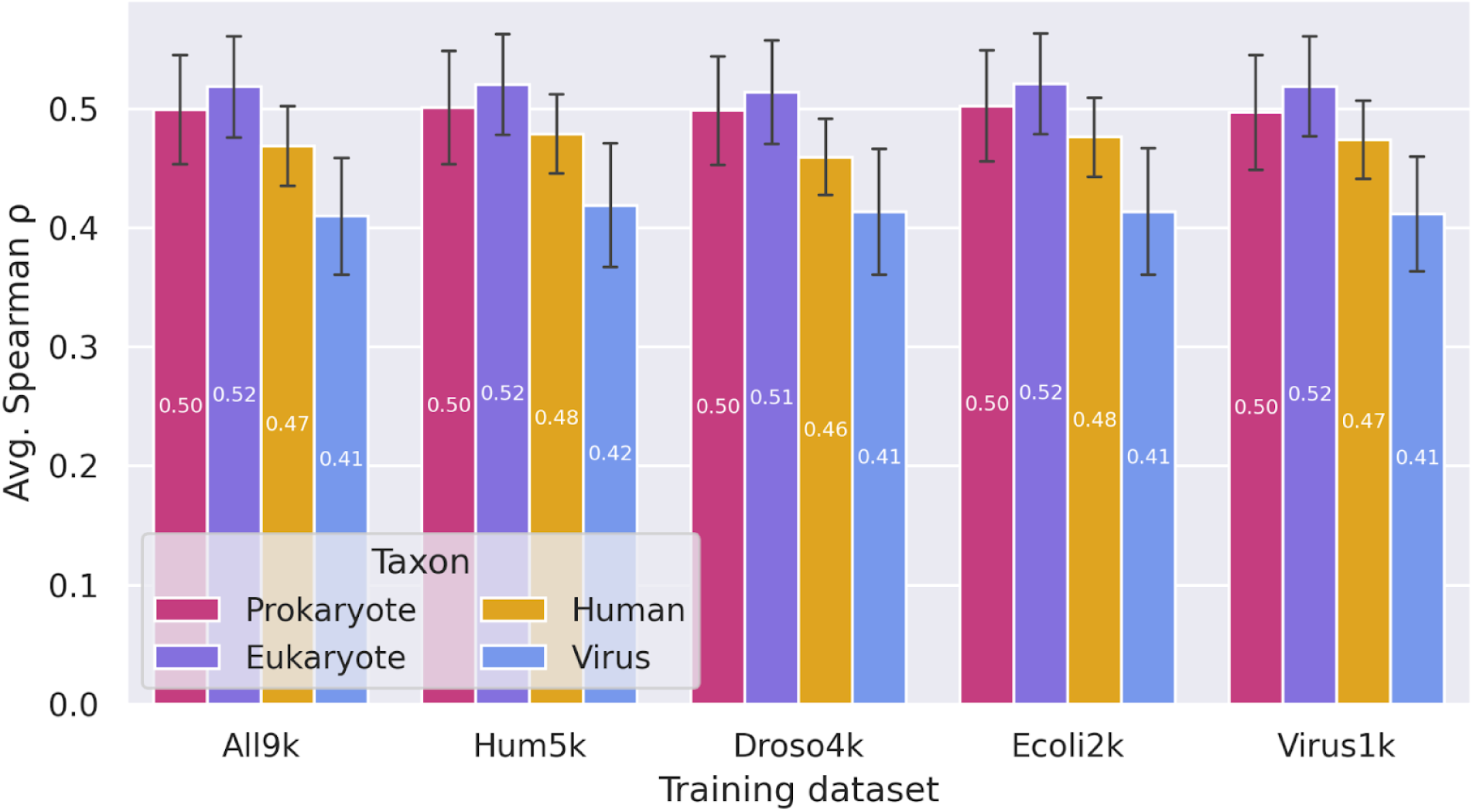
Per-taxon performance of VespaG independent of taxa included in training. For each training set, indicated on the x-axis, we reported the average Spearman correlations computed for each of the five taxa represented in ProteinGym benchmark (Notin et al. 2023). The error bars show the 95% confidence interval. Regardless of training data (all data sets were redundancy reduced, names reflect training and bars the test set; *All9k*: about 9,000 proteins mixing all taxa shown to the right; *Hum5k*: ∼9,000 human proteins, *Droso4k*: ∼4,000 fruit fly proteins (*drosophila melanogaster*) , *Ecoli2k*: ∼2,000 E.coli proteins (*Escherichia coli*), *Virus1k*: mix of ∼1,500 viral proteins), VespaG generalized equally well for all taxa assessed. For all training sets, it performed best for prokaryotes and non-human eukaryotes, followed by human. Even when trained explicitly on viral proteins, VespaG performed the worst for viral proteins.

### VespaG predictions blazingly fast

Out of the top performing methods evaluated on the ProteinGym benchmark, VespaG was, in our hands, the most scalable for proteome-wide analyses. Inference on CPU with VespaG needed 5.7 seconds (s) for the 73 unique proteins from ProteinGym first iteration (SOM Figure S4) on low-end hardware (Intel i7-1355U with 12x5 GHz, 1.3 GB RAM, no GPU; SOM Table S4). GEMME completed the predictions on the same hardware in 1.27 hours (h) (4.2 GB RAM, SOM Table S4). Even when considering the time required for input pre-processing, the highly efficient MSA generation of ColabFold could not overcome the runtime advantage of VespaG with a total runtime <1h on a consumer CPU (SOM Table S5). Accessing high-end GPU and CPU resources for pre-processing led to a total execution time of ∼1 minute (min) (64.3s) for VespaG versus ∼90 min (5,468.4s) for GEMME (SOM Table S4-5). By comparison, computing zero-shot ESM-2 log-odds took ∼5.35 days and VESPA required 17h (SOM Table S4).

Thus, VespaG was five orders of magnitude faster than ESM-2 (factor 10^5, i.e., 100,000-times), and three orders of magnitude faster than GEMME and VESPA. The authors of PoET observed their method and TranceptEVE L to be three orders of magnitude slower than GEMME (Truong Jr and Bepler 2023) providing some base to triangulate an estimate between those and VespaG (∼ factor of 10^6 faster). Additionally, VespaG, unlike other pLM-based methods, does not require a GPU for fast inference (SOM Table S4). On a consumer-grade laptop (Methods), VespaG computed the entire single-site mutational landscape for a human proteome with 20k proteins (Sinitcyn et al. 2023) in fewer than 30 minutes. On the same machine, at the same time, GEMME completed predictions for 25 proteins. If we assumed that methods such as PoET or TranceptEVE L were executable on low-end hardware, they would have processed 0.025 proteins.

## Discussion

In this work, we explored the possibility of modeling the sequence-phenotype relationship by learning a simple mapping function from protein Language Model (pLM) representations, or embeddings, to evolutionary scores predicted by an expert method. We demonstrated the validity of this approach on several hundred diverse proteins across different organisms. The performance of the resulting method, dubbed VespaG, reached that of much more sophisticated methods.

Using predicted scores instead of curating a dataset of experimental measures allowed the creation of a larger training set (totaling 39M mutations) than those used previously. By comparison, the SNAP2 development set contained about 100,000 mutational effect annotations from 10,000 proteins (Hecht, Bromberg, and Rost 2015). In addition, exploiting only the pLM embedding of the wild-type protein of interest, instead of explicitly modeling its mutants through log-odds probability estimates, enabled reaching a much higher efficiency and inference speed than previous pLM-based methods. The feasibility of this strategy also emphasized the usefulness of the information encoded in a wild-type protein query embedding for assessing all its variants.

### VespaG reached SOTA despite its simplicity

The fact that prediction accuracy for VespaG reached the state-of-the-art (SOTA) level proves that even relatively shallow neural networks (660k free parameters) can effectively leverage the knowledge encoded in an unsupervised method such as GEMME (Laine, Karami, and Carbone 2019). The transparency of the GEMME algorithm and the simplicity of VespaG architecture guarantee some level of interpretability of the predictions, which is highly valuable for investigating the mechanisms underlying mutational outcomes.

A fundamental difference between student (VespaG) and teacher (GEMME) is its usage of a universal protein representation space. More specifically, VespaG can relate proteins with each other via representations generated by a pLM pre-trained over a huge diversity of natural protein sequences across protein families. This property allows VespaG to generalize across organisms without considering any specific input generation or training schema. Training VespaG on a few thousand proteins from either *Homo sapiens*, or *Drosophila melanogaster*, or *Escherichia coli* sufficed to produce high quality predictions on a diverse set of proteins. For eukaryotic and prokaryotic proteins, the pLM-based student VespaG performed overall numerically higher and more consistently than the MSA-based teacher GEMME. For instance, VespaG improved for cases such as ubiquitin which do not follow the general trends between evolutionary conservation and mutational outcomes.

Nevertheless, biases in the pLM representation space may lead to poor predictions for some protein families. Namely, the ProteinGym assessment consistently reported lower accuracy on viral proteins for all zero-shot pLM predictors (Notin et al. 2023). Our results demonstrated that supervising on GEMME scores partially counterbalanced this trend. VespaG’s performance decreased for viral proteins, even when explicitly trained on those, but it performed favorably compared to the pLMs ESM-2 and SaProt. The gap in performance between the student, VespaG, and the teacher, GEMME, for this class of proteins, suggests specific properties of the embeddings or the evolutionary scores preventing an accurate mapping between them. Despite retaining high accuracy, GEMME evolutionary scores for viral proteins have a lower resolution than for organismal proteins (SOM Fig. S2). Many mutations are assigned the same score, likely reflecting the comparatively lower variability of the associated input MSAs (Table 1). Nevertheless, the fact that VespaG trained exclusively on viral proteins exhibits a high predictive capability on organismal proteins (Fig. 4) suggests the impact of this resolution loss is limited. It further supports the hypothesis that the embeddings computed by the pre-trained pLMs for viral proteins are intrinsically noisy. Possibly, viral proteins are simply too under-represented in the training of pLMs, due to a comparatively small number and low diversity (Ding and Steinhardt 2024; Elnaggar et al. 2021; Lin et al. 2023; The UniProt Consortium et al. 2023). In addition, the pLMs may struggle to capture the inherent peculiarities of viral protein evolution (Koonin, Dolja, and Krupovic 2022). Structurally and functionally relevant evolutionary constraints are expected to manifest through smaller differences in viral protein sequences compared to other taxa, warranting a special treatment of these sequences for extracting co-variations (Hopf et al. 2017). A future improvement of pLMs could be to develop viral-specific fine-tuning steps.

Finally, we showed that VespaG’s independence from alignments combined with its inherent generalizability enables tackling *de novo* designed proteins. Most established mutation effect prediction methods, such as GEMME, largely succeed due to evolutionary information derived from MSAs and tend to barely outperform random for single sequences (Hecht, Bromberg, and Rost 2015). Given the absence of reference data for *de novo* proteins, MSA-based tools often fail to provide any result. At the same time, pLMs such as ESM-2 tend to provide unreliable estimates of their respective properties. Although we observed a drop in performance compared to natural proteins, we can envision using VespaG for fast screens before applying future methods adapted to that problem or as a guide for designing more biocompatible *de novo* proteins.

### Saving resources as criterion

Although we acknowledge the interest of the pairwise comparison-based predictor ranking scheme introduced recently ((Atlas of Variant Effects Alliance n.d.; Livesey and Marsh 2023)), we decided to keep the analysis simpler, tuned to the perspective of the ProteinGym benchmark. Our motivation for this choice is that ranks for individual methods remain short lived although trends for the field appear more stable ((Atlas of Variant Effects Alliance n.d.; Livesey and Marsh 2023); Notin et al. 2023).

Beyond providing a proof-of-principle for the success of teacher-student strategy in the field of variant effect prediction, our work emphasises the possibility to improve speed and reduction of energy consumption. VespaG provides an important example for an extremely fast, cost-efficient, simple tool that invites saving resources at very little cost in terms of accuracy. These properties are highly valuable in a context where many researchers are interested in variant effect predictions for proteomes for which no data is available. Moreover, VespaG’s simplicity stands out in the environment of SOTA predictors. Looking at, *e.g.*, the ablation study of AlphaMissense (Cheng et al. 2023), we note how many impressively complex aspects of the method make that method reach its top-level performance. VespaG reaches a similar level without any of that: not using complex machine learning on the side of learning from the teacher, no 3D structure, no MSA, no minor allele frequency-based loss function, no database distillation, asf.

Being orders of magnitudes faster than prior methods, VespaG makes it possible, for instance, to explore the effect of a mutation arising in many different contexts such as protein engineering, opening the way to a systematic assessment of epistasis. Additionally, the student-teacher setup of VespaG is easily adaptable to novel pLM input and additional features.

### Gain of speed at the expense of interpretability?

GEMME reaches its SOTA-level performance by optimizing only two simple parameters: the conservation of a position in a family of related proteins and the distance of a variant on the tree. Simply put, GEMME predicts effect when variants deviate from the observed conservation pattern and neutral when the variants have been observed close on the tree. *Per se*, VespaG has no such interpretability. As it learned from GEMME, users can replace “strongly predicted effect” to imply variant against conservation even without seeing the MSA, and conversely “strongly predicted neutral” as examples observed nearby on the tree. In fact, in contrast to GEMME, VespaG quantifies the strength of the prediction. This in itself seems an important feature relevant for users. For the analysis of particular variants, users might want to actually generate MSAs and trees on their own to support their rationales. However, neither any of the two, nor - to the best of our knowledge - any of the other SOTA methods, directly generate a hypothesis for how a variant may disrupt the details of molecular function.

### Interpretable models of variant effects?

For some particular applications, users might be interested to have the “molecular” or “mechanical” explanation why a particular variant affects protein function or why it is neutral. Future efforts may aim at more explicitly encoding evolutionary semantics in the pLM representation space, e.g., by exploiting GEMME intermediate results (evolutionary distances). Modeling and tracing the evolutionary history of natural sequences in that space could provide the key to pinpoint gain-of-function mutations and distinguish them from loss-of-function ones, a major current challenge for the field. Another direction for future improvements could be the combination of effect prediction methods, such as GEMME, or VespaG, with methods optimized to predict certain phenotypes as has been proposed before (Ioannidis et al. 2016). For instance, many methods predicting thermodynamic stability changes (Benevenuta et al. 2021; Chen et al. 2020; Li et al. 2020) exploit complementary information about 3D structures, and the expert-guided ML method RaSP already enables large-scale screens (Blaabjerg et al. 2023). Ideally, the field would advance by adding semi-automated tools to turn predictions into mechanical explanations using annotations and predictions of protein structure and function.

In conclusion, VespaG closes the gap in performance between the best and the fastest missense amino acid variant effect predictors. For an unprecedentedly small trade-off in performance, it can predict variants several orders of magnitude faster than other state-of-the-art methods.

## Methods

### Related work

A key component of computational variant effect predictors is the ability to capture arbitrary range dependencies between amino acid residues. A majority of predictors extracts these dependencies from an input multiple sequence alignment (MSA) generated from large protein sequence databases. Some rely on the statistical inference of pairwise couplings (Figliuzzi et al. 2016; Hopf et al. 2017), others on the implicit account for global context with latent variables (Frazer et al. 2021; Riesselman, Ingraham, and Marks 2018). They remain computationally costly due to the large number of inferred parameters and strongly depend on the input alignment’s variability.

More recently, using representations generated by protein language models (pLMs) emerged as an alternative to using MSAs as input (Brandes et al. 2023; Cheng et al. 2023; Marquet et al. 2022; Meier et al. 2021; Nijkamp et al. 2022; Notin, Dias, et al. 2022). These high-capacity pLM transformer architectures, borrowed from natural language processing, learn to reconstruct masked or missing amino acids in an input query sequence (Elnaggar et al. 2021; Lin et al. 2023). They model raw protein sequence data over large databases, thereby capturing evolutionary constraints that generalize across protein families. Once trained, they can serve as zero-shot variant effect predictors by estimating the likelihood of each amino acid at each position.

A limitation of pLMs is that they overlook natural protein sequences’ evolutionary history and relationships. Strategies for overcoming this oversimplification aim at encoding evolutionary semantics in the pLMs representation space, *e.g.*, by augmenting the input with a multiple sequence alignment (Rao et al. 2021). The latter informs the model about sequences evolutionary related to the input query and how their amino acids match. The integration of weak labels coming from inter-individual polymorphisms and of a physical prior through supervised learning of the 3D structure, as AlphaMissense does, further enhances the predictive performances (Cheng et al. 2023). Alternatively, the predictor VESPA combines the pLM-derived log-odds substitution scores with evolutionary conservation levels predicted from the learned protein representations (Marquet et al. 2022). The predictors Tranception (Notin, Dias, et al. 2022) and PoET (Truong Jr and Bepler 2023) have also explored retrieval-augmented strategies by conditioning the predictions of a pre-trained pLM on a set of related raw or aligned protein sequences at inference time. Others successfully included structural information to enhance performance for various supervised and unsupervised downstream prediction tasks (Heinzinger et al. 2024; Su et al. 2024; Tan et al. 2024).

### Comparison to State-of-the-art methods

We chose to compare our methods to the following seven state-of-the-art methods on the ProteinGym test set. *GEMME* (Laine, Karami, and Carbone 2019) as it is (1) tied for the best performing method on ProteinGym (Notin et al. 2023), (2) a purely MSA-based method not using machine learning, and (3) was used to annotate VespaG’s training data. Zero-shot log-odds by the pLMs *ESM-2* (Lin et al. 2023), the sequence-only pLM used as input to VespaG (for details see *Model Specifications - Input features*), and *SaProt* (Su et al. 2024), a top-ranked pLM in ProteinGym which takes structure and sequence as input. *TranceptEVE L* (Notin, Niekerk, et al. 2022) as it is the best performing method on ProteinGym next to GEMME and SaProt, and because it is a hybrid model, making use of both MSAs and pLM embeddings as input, combining the previously developed autoregressive Tranception (Notin, Dias, et al. 2022) with the Bayesian variational autoencoder EVE (Frazer et al. 2021). *PoET* (Truong Jr and Bepler 2023), a recently developed autoregressive generative method modeling protein families as sequences-of-sequences and slightly outperforming other methods against the first version of the ProteinGym set (although not currently included in the benchmark). *AlphaMissense* (Cheng et al. 2023), also recently introduced and building up on the protein structure predictor AlphaFold (Jumper et al. 2021) by incorporating population frequency data. Similar to PoET, AlphaMissense was evaluated on the first ProteinGym iteration but is not included in the updated benchmark. Lastly, we compare to *VESPA* (Marquet et al. 2022), the currently best purely sequence pLM-based method in ProteinGym. VESPA takes pLM embeddings as input to predict per-residue conservation scores and combines them with per-mutation protein-dependent log-odds scores and per-mutation protein-independent substitution scores.

We computed VESPA and GEMME predictions for both test sets, while we downloaded TranceptEVE L, SaProt and ESM-2 predictions from ProteinGym. For PoET, the authors provided Spearman correlation values for each experimental assay in the first iteration of the ProteinGym test set on request. For AlphaMissense, we extracted pre-computed per-assay scores of the first ProteinGym iteration from the supplemental materials.

Due to the different ways of accessing or computing the predictions of GEMME, TranceptEVE, PoET, and AlphaMissense, we did not evaluate the effect of using different MSAs as input. GEMME predictions were generated through the MMseqs2-based ColabFold protocol as described in (Abakarova et al. 2023). Further, we did not benchmark ensemble methods such as PoET+GEMME (Truong Jr and Bepler 2023) on the test sets as ensembling any other method with VespaG would eradicate its speed advantage. To assess thermodynamic folding stability of de novo domains, we only compared VespaG with ESM-2 and GEMME, since precomputed results were not available for the other methods.

### Method development

#### Datasets

In the following, the datasets applied in this work are described. We utilized ten different datasets, one based on the human proteome (Hum5k) for training and validation, and four re-training datasets (Droso4k, Ecoli2k, Virus1k, All9k). All training data was annotated using evolutionary-informed substitution scores computed by GEMME. Additionally, we compared VespaG against SOTA methods on the two test sets *ProteinGym* (with nine subsets) and *StabilityDeNovo146*.

#### Test data

##### ProteinGym Substitution Benchmark

The substitution benchmark ProteinGym (Notin et al. 2023) comprised 217 deep mutational scans (DMS) from 187 unique proteins with diverse lengths (37 - 3,423 residues with a median of 245), protein families (*e.g.*, polymerases, tumor suppressors, kinases, transcription factors), sizes, functions (e.g. drug resistance, ligand binding, viral replication, thermostability), and taxa, totalling about 696k single missense variants and 1.76M multiple missense mutations from 69 of the 217 proteins (SOM Fig. S2a). Protein sequences and experimental scores were accessed from the ProteinGym GitHub repository (Notin 2024). The ProteinGym substitution benchmark was not redundancy reduced, and we divided the ProteinGym into subsets based on organisms, function, and time of availability. The first split by organism and can contain any function: (1) *PGOrganismal189* with 189 assays on 161 prokaryotic and eukaryotic proteins, and (2) *PGViral28* with 28 assays on 26 viral proteins. The second split was based on function and can contain prokaryotic, eukaryotic and viral proteins: (3) *PGStability66* with 66 assays assessing stability, (4) *PGActivity43* with with 43 assays assessing activity, (5) *PGBinding13* with 13 assays assessing binding, (6) *PGExpression18* with 18 assays assessing organismal expression, and (7) *PGFitness77* with 77 assays assessing fitness. The last splits were added to assess methods for which predictions were available only for the first iteration of ProteinGym. This first iteration contained 87 DMS from 73 unique proteins (72 - 3,423 residues with a median of 379), totalling about 1.5M variants with mostly single and, for 11 proteins, multiple missense mutations (SOM Fig. S4b). The set was divided into (8) *PGOrganismal66,* containing 66 assays of 54 proteins with 1.4M variants, and (9) *PGViral21*, containing 21 assays of 19 proteins with 184K variants.

##### StabilityDeNovo146

The dataset by Tsuboyama et al. 2023 includes DMS measuring stability under identical conditions of all single substitutions, deletions and two insertions in 266 native and 146 *de novo* designed domains (40–72 residues). This is the most comprehensive available dataset assessing how amino acid substitutions affect thermodynamic folding stability. The ProteinGym substitution benchmark includes 64 native proteins from this dataset (all included in *PGStability66*). To additionally assess the predictors on *de novo* domains, we created the test set *StabilityDeNovo146* with 123k variants across 146 unique proteins designed using TrRosetta (Yang et al. 2020) with the hallucination protocol described in (Anishchenko et al. 2021; Norn et al. 2021) or the blueprint-based approach described in (Huang et al. 2011; Kim et al. 2022). We selected all mutations with tabulated free energy changes (ΔΔG) annotations, discarded all deletions, insertions, and wild-type sequences, and averaged multiple measurements for the same mutation.

#### Training data

To generate training data, we constructed a main set based on the *Homo sapiens* proteome and additional sets representing a diverse set of origins. These included proteomes from *Drosophila melanogaster* and *Escherichia coli,* as well as all viral proteins included in Swiss-Prot (Table 1). Each training dataset was curated following the same process of first downloading the UniProt (The UniProt Consortium et al. 2023) reference proteome(s) with one protein sequence per gene for the respective organism or set of organisms and removing any proteins with a sequence length smaller than or equal to 25 residues or longer than or equal to 1024. The dataset sizes are presented in Table 1.

We redundancy reduced the training data in two steps, firstly against the test data to prevent data leakage and secondly against themselves to reduce the number of training samples. To do so, we clustered the protein sequences using UniqueProt (Mika 2003) with an HSSP (homology-derived secondary structure of proteins)-value < 0, corresponding to no pair of proteins in the redundancy reduced data set having over 20% pairwise sequence identity over 250 aligned residues (Rost 1999; Sander and Schneider 1991). This criterion is stricter than the classically used threshold of 30% overall sequence identity. It accounts for the fact that protein structures can show high similarity even at lower sequence similarity levels (Rost 1999). To improve runtime, we modified the original UniqueProt protocol (Olenyi, Tobias et al. n.d.) by replacing BLAST (Altschul et al. 1997) with MMseqs2 (Steinegger and Söding 2017). Additionally, we discarded alignments of fewer than 50 residues for pairs of sequences with more than 180 residues as they provide only a relatively weak support for similarity. Finally, we generated training and validation sets using a random 80/20 split.

To circumvent the need for a large, comprehensive set of experimental variant effect annotations, we employed the established method GEMME (see Evaluation/SOTA Methods) following the protocol introduced in (Abakarova et al. 2023). Specifically, for each protein from the training set, we retrieved and aligned a set of homologous sequences with the MMseqs2-based multiple sequence alignment (MSA) generation strategy implemented in ColabFold (Mirdita et al. 2022). We then used the generated MSA as input for GEMME. GEMME outputs a complete substitution matrix of dimension L x 20, with L being the length (in residue) of the input query protein sequence. GEMME scores range from -10 to 2. Drawing from our previous findings (Abakarova et al. 2023), we flagged the mutational landscapes derived from fewer than a couple hundred homologous sequences as lowly confident.

### Model Specifications

#### Input features

All developed models rely solely on embeddings computed from pre-trained pLMs as input. Specifically, we used ProtT5-XL-U50 (Elnaggar et al. 2021), an encoder-decoder transformer architecture trained on the Big Fantastic Database (Steinegger and Söding 2017) and fine-tuned on UniRef50, and ESM2-T36-3B-UR50 (Lin et al. 2023), a BERT (Devlin et al. 2019) style 3-billion-parameter encoder-only transformer architecture trained on all clusters from Uniref50, augmented by sampling sequences from the Uniref90 clusters of the representative chains (excluding artificial sequences). In the following, we refer to these pLMs as ProtT5 and ESM-2, respectively. For both pLMs, we downloaded the encoder weights from HuggingFace (Wolf et al. 2020) and extracted the embeddings from the encoder’s last hidden layer. These embeddings comprise 1024-dimensional vectors for each residue in a sequence for ProtT5 (Elnaggar et al. 2021) and 2560-dimensional vectors for ESM-2 (Liu and Carrigan 2024). There is no length restriction for either pLM at inference, so proteins were processed in full. A guide to embedding extraction for ProtT5 and ESM-2 can also be found in our GitHub repository https://github.com/jschlensok/vespag. We used the pre-trained pLMs as is, without fine-tuning their weights, and without combining the embeddings either by concatenating the input or averaging the outputs.

#### Configuration

We built the predictors with the following architectures: (1) Linear regression, *i.e.*, a feed-forward neural network (FNN, (LeCun, Bengio, and Hinton 2015)) without any hidden layer, dubbed *LinReg*; (2) FNN with one dense hidden layer, called *VespaG*; (3) FNN with two hidden layers, called *FNN_2_layer*; (4) Convolutional neural network (CNN, (LeCun, Bengio, and Hinton 2015)) with one 1-dimensional convolution and two hidden dense layers, referred to as *CNN*; and (5) an ensemble of separately optimized FNN and CNN (with the same architecture as the best stand-alone model for each architecture), with the output being the mean of the two networks.

All layers were linked through the LeakyReLu activation function (Agarap 2019; Xu et al. 2015), as well as dropout (Srivastava et al. 2014). No transformation was applied to the output of the final layer. We implemented the models in PyTorch (Paszke et al. 2019) v1.13.1 using Python 3.10.8. They were trained using the AdamW (Kingma and Ba 2017; Loshchilov and Hutter 2019) optimizer with an initial learning rate of 10e-4, decaying with a decay factor of 0.33 and a patience of 8 epochs using mean squared error (MSE) loss between predicted and target GEMME scores. Each batch contained 25K residues. We applied early stopping based on the validation partition loss, with a patience of 10 epochs. The maximal training duration was 200 epochs. Hyperparameters were optimized through exhaustive parameter search on the validation split of the *Hum5k* dataset for each architecture. The same hyperparameters were then used to re-train separate models on each of the additional datasets (Droso4k, Ecoli2k, Virus1k, and All9k), using the same training scheme (SOM Table S1). The training dataset and hyperparameters of the final model were selected based on validation loss.

### Evaluation

#### Performance measures

To assess the performance of the predictors, we relied on the Spearman rank correlation coefficient: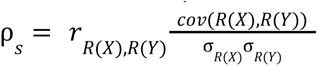, with raw scores 𝑋, 𝑌 as ranks 𝑅(𝑋), 𝑅(𝑌), Pearson correlation coefficient *r*_𝑅(𝑋),𝑅(𝑌)_, covariance 𝑐𝑜𝑣(𝑅(𝑋), 𝑅(𝑌)), and standard deviations σ_𝑅(𝑋)_, σ_𝑅(𝑌)_. This metric enables quantifying the strength of non-linear relationships between predicted and experimental scores. The Spearman correlation coefficient was computed for each DMS experiment separately. For proteins covered by multiple DMS experiments, we averaged the correlation over the experiments before computing the mean over all proteins.

#### Error estimates

To assess the significance of the performance differences between the evaluated predictors, we computed symmetric 95% confidence intervals (CI) µ ± 1. 96 * 𝑆𝐸𝑀, with 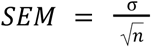 as the Standard Error of the Mean over n = 1,000 bootstraps using sampling with replacement from the respective datasets. The terms µ and σ are the mean and standard deviation of the bootstrap distribution, respectively. In addition, we performed one-tailed paired t-tests on the distributions of Spearman rank correlation coefficients. For any pair of methods, the test statistic is expressed as, 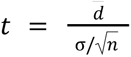 where *d̄* and σ are the mean and standard deviation of the 𝑛 correlation coefficient differences. The associated p-value reflects the probability of observing the test statistic under the null hypothesis (no significant difference, the true mean of differences is zero). It is obtained by comparing 𝑡 to a 𝑡-distribution with 𝑛-1 degrees of freedom.

#### Baseline

To compare model performances on the validation set to an untrained random baseline retaining on the target distribution, we randomly permuted the GEMME target scores for each protein, i.e. randomly shuffled all values of the Lx20 GEMME output matrix.

#### Multi-mutations

To score combinations of multiple substitutions, we add the scores of their constituent single substitutions, a technique introduced previously (Meier et al. 2021). For instance, in the case of the double mutation *M1A:G2N*, i.e. the mutation of a Methionine to an Alanine at the first residue of the protein and a Glycine to an Asparagine at the second, the score of the double mutation is the sum of the computed scores of the single mutations *M1A* and *G2N*.

#### Runtime

We measured the wall-clock runtime of VESPA, GEMME, and VespaG for 1.6M mutations of the 73 unique proteins of the first iteration of the ProteinGym test set using 32 CPU cores of an Intel Xeon Gold 6248 at 2.50 GHz and 64 GB of DDR4 ECC RAM. VESPA, in addition, was allotted an Nvidia Quadro RTX 8000 GPU with 48 GB VRAM. Runtimes for other methods could not be obtained due to computational limitations. We did not measure the runtime for the generation of the input MSAs for GEMME, nor the input embeddings for the embedding-based methods since we used pre-computed data.

#### Score Normalization

To ease score interpretability, VespaG scores are normalized to the [0, 1] interval by applying the logistic function 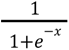 to the raw GEMME scores. This behavior can be toggled off by the user if desired.

## Declarations

## Supporting information

SOM

## Acknowledgements

This work was granted access to the high-performance computing resources of the SACADO MeSU platform at Sorbonne Université. Thanks to Nikita Kugut for invaluable help with administrative aspects of this work, as well as Tim Karl and Tobias Olenyi for technical aspects. Thanks to the group around Pascal Notin for creating the benchmark ProteinGym, and to all developers who make their methods freely available. Thanks to all experimentalists for making their data available - with special thanks to the group around Gabriel J. Rocklin for developing the mega-scale experimental analysis of protein folding stability.

## Data and Code Availability

The source code and model weights of this work are freely available at https://github.com/JSchlensok/VespaG. The data used for development of VespaG and VespaG predictions are freely available at https://doi.org/10.5281/zenodo.11085958.

## Author Contributions

C.M. supervised the method implementation and evaluation, and co-wrote the manuscript. J.S. implemented and evaluated the method and wrote the initial draft. M.A. computed GEMME scores and assisted in the usage of the tool. B.R. co-conceived and guided the work. E.L. supervised the work and co-wrote the manuscript. J.S., C.M., M.A and E.L. produced and analyzed the results. All authors edited, read, and approved the final manuscript.

