## Supplementary material for "Expert-guided protein Language Models enable accurate and blazingly fast fitness prediction": SOM

### Description

This supporting material provides information to help the reader with reproducing and re-interpreting the results of the main article on VespaG, a blazingly fast amino acid variant effect predictor, leveraging embeddings of protein language models (pLMs) as input to a minimal deep learning model. Table S1 and Figures S1 and S2 highlight method development w.r.t. hyperparameter configurations and pLM types. Figures S3 and S4 show test set properties. Tables S2-S3 and Figures S3-S8 present more details on the Spearman correlation between experimental results and evaluated methods. Tables S4 and S5 highlight runtime performance of evaluated methods.

**TABLE OF CONTENTS**

1. Table S1: Hyperparameters and number of free parameters.
2. Table S2: Spearman Correlation ρ between ProteinGym substitution benchmark and SOTA methods.
3. Table S3: Spearman Correlation ρ with ProteinGym substitution benchmark for methods grouped by number of point mutations in mutants.
4. Table S4: Runtime comparison for VespaG, GEMME, VESPA, and ESM-2 (3B) inference.
5. Table S5: Runtime comparison for VespaG, GEMME, and VESPA pre-processing steps.
6. Figure S1: Influence of the architecture and the input embeddings on VespaG's validation loss.
7. Figure S2: Influence of the training dataset and the input embeddings on VespaG's validation loss.
8. Figure S3: Properties of the ProteinGym substitution benchmark set.
9. Figure S4: Properties of the first iteration of the ProteinGym substitution benchmark set.
10. Figure S5: Average Spearman correlation of VespaG and SOTA methods on unique proteins of *ProteinGym217.*
11. Figure S6: Consensuality of the predictions on ProteinGymOrganismal189.
12. Figure S7: Average Spearman correlation of VespaG and SOTA methods on the first iteration of ProteinGym.
13. Figure S8: Average Spearman correlation of VespaG and SOTA methods on 189 organismal DMS assays of the ProteinGym benchmark.
14. Figure S9: Average Spearman correlation of VespaG and SOTA methods on 28 viral DMS assays of the ProteinGym benchmark.
15. Figure S10: Comparison of Spearman correlation depending on the input alignment.
16. Figure S11: Average Spearman correlation between predicted mutational effect scores and experimental ΔΔG scores.
17. References

| Model | Best hyperparameters | # free parameters (ProtT5/ESM-2)* |
|---|---|---|
| LinReg | Dropout of 0.4 | 1025 / 2561 |
| **FNN_1_layer (*VespaG*)** | Hidden layer with size 256, dropout 0.2. | 267k / 660k |
| FNN_2_layer | Hidden layers of sizes 256 and 64 without dropout. | 280k / 673k |
| CNN | 1D convolution from input to 256 channels with kernel size 7 and padding 3 with dropout rate 0.2. Fully-connected hidden layers of size 256 and 64 without dropout. | 1.91m / 4.67m |
| FNN+CNN mean | FNN_2_layer + CNN | 2.19m / 5.34m |

**Table S1**: **Hyperparameters and number of free parameters.** We built the predictors with 5 architectures, for each the selected hyperparameters and number of free parameters is listed. Hyperparameters were optimized through exhaustive parameter search on the validation split of the *Hum5k* dataset for each architecture. Methods are: (1) Linear regression, *i.e.*, a feed-forward neural network (FNN) without any hidden layer, dubbed *LinReg*; (2) FNN with one hidden layer, called *FNN_1_layer (VespaG)*; (3) FNN with two hidden layers, called *FNN_2_layer*; (4) Convolutional neural network (CNN) with one 1-dimensional convolution and two hidden dense layers, referred to as *CNN*; and (5) an ensemble of separately optimized FNN and CNN (with the same architecture as the best stand-alone model for each architecture), with the output being the mean of the two networks.
*Size of input embeddings: ProtT5 1024xL and ESM-2 2560xL

| ProteinGym Subset | VespaG | GEMME | TranceptEVE L | VESPA | ESM-2 3B | SaProt 650M |
|---|---|---|---|---|---|---|
| *PG217* (averaged per-protein) | 0.480 ±0.021 | **0.486 ±0.021** | 0.474 ±0.021 | 0.460 ±0.02 0 | 0.430 ±0.028 | 0.472 ±0.027 |
| *PG217* (weighted average per-function) | 0.459 ±0.050 | **0.464 ±0.041** | 0.456 ±0.040 | 0.436 ±0.043 | 0.406 ±0.057 | 0.457 ±0.074 |
| *PGOrganismal189* | 0.491 ±0.024 | 0.490 ±0.024 | 0.478 ±0.023 | 0.464 ±0.023 | 0.455 ±0.027 | **0.500 ±0.026** |
| *PGViral28* | 0.414 ±0.049 | **0.462 ±0.045** | 0.453 ±0.052 | 0.432 ±0.045 | 0.274 ±0.087 | 0.300 ±0.083 |
| *PGActivity43* | **0.494 ±0.043** | 0.493 ±0.045 | 0.487 ±0.045 | 0.468 ±0.043 | 0.417 ±0.061 | 0.458 ±0.055 |
| *PGBinding13* | 0.370 ±0.086 | **0.397 ±0.090** | 0.376 ±0.085 | 0.366 ±0.076 | 0.321 ±0.103 | 0.379 ±0.075 |
| *PGExpression18* | 0.456 ±0.063 | 0.443 ±0.066 | 0.457 ±0.065 | 0.404 ±0.067 | 0.403 ±0.081 | **0.488 ±0.064** |
| *PGFitness77* | 0.441 ±0.036 | **0.460 ±0.034** | **0.460 ±0.034** | 0.440 ±0.035 | 0.379 ±0.048 | 0.367 ±0.046 |
| *PGStability66* | 0.533 ±0.036 | 0.528 ±0.038 | 0.500 ±0.040 | 0.500 ±0.037 | 0.509 ±0.040 | **0.592 ±0.033** |

**Table S2**: Spearman correlation coefficient $\rho$ between predicted and experimental substitution effect scores on ProteinGym substitution benchmark (with several subsets) for methods VespaG, GEMME (Laine et al., 2019), TranceptEVE L (Notin et al., 2022), VESPA (Marquet et al., 2022), ESM-2 (3B) (Lin et al., 2023), and SaProt (650M) (Su et al., 2024). Cells show mean $\rho$ ± standard error for subsets: *ProteinGym217 (per-protein)* - all 217 DMS, weighted by the number of DMS per protein, *ProteinGym217 (per-function)* - all 217 DMS, weighted by the number of DMS per protein and the number of assays per category. All others are weighted by the number of DMS per protein: *ProteinGymOrganismal189* - 189 eukaryotic and prokaryotic DMS, *ProteinGymViral28* - 28 viral DMS, *ProteinGymActivity43* - 43 DMS on activity, *ProteinGymBinding13* - 13 DMS on binding, *ProteinGymExpression18* - 18 DMS on expression, *ProteinGymFitness77* - 77 DMS on organismal fitness, and *ProteinGymStability66* - 66 DMS on stability. Numerically highest values per row highlighted in bold. Standard error bootstrapped over unique proteins for all subsets except *ProteinGym217 (per-function)* where it was instead bootstrapped over the 5 functional categories.

| Mutational Depth | Distribution | | Avg. Spearman ρ | | | | |
|---|---|---|---|---|---|---|---|
| | # assays | # mutations | VespaG | GEMME | Trancept EVE L | VESPA | ESM-2 (3B) |
| 1 | 217 | 696,311 | 0.462 | **0.464** | 0.391 | 0.396 | 0.366 |
| 2 | 69 | 826,245 | 0.249 | **0.292** | 0.256 | 0.183 | 0.208 |
| 3 | 11 | 84,134 | 0.347 | **0.376** | 0.250 | 0.324 | 0.179 |
| 4 | 11 | 187,850 | 0.319 | **0.349** | 0.211 | 0.278 | 0.150 |
| 5+ | 9 | 671,227 | 0.367 | **0.423** | 0.249 | 0.287 | 0.185 |

**Table S3**: **Spearman correlation coefficient ρ between predicted and experimental substitution effect scores on ProteinGym substitution benchmark partitioned by mutational depth.** We report results for methods VespaG, GEMME (Laine et al., 2019), TranceptEVE L (Notin et al., 2022), VESPA (Marquet et al., 2022), and ESM-2 (3B) (Lin et al., 2023). Numerically highest values per row highlighted in bold. Results were not averaged by protein or assay type. No per-mutant predictions for SaProt were available to download via ProteinGym as of June 2024.

| Inference | | | |
|---|---|---|---|
| **Tool** | **Hardware** | **Runtime [s]** | **Memory usage** |
| VespaG | Consumer CPU | 5.7 | 1.3 GB |
| GEMME | Consumer CPU | 4,561.4 | 4.2 GB |
| ESM-2 (3B) log-odds ratios | High-end GPU | 462,417.34 | 48 GB |
| VESPA | High-end GPU | 63,693.0 | 48 GB |

**Table S4: Runtime of VespaG, GEMME, ESM-2 (3B) and VESPA inference on 73 unique proteins of the first iteration of the ProteinGym substitution benchmark (Notin et al., 2023) with precomputed input.** Hardware used: consumer CPU - Intel i7-1355U with 12x5 GHz, high-end GPU - Nvidia Quattro RTX 8000 or Nvidia RTX 6000 (both 48GB VRAM). Methods benchmarked: VespaG with pre-computed ESM-2 (3B) per-residue embeddings, GEMME with pre-computed ColabFold alignments (Laine et al., 2019; Mirdita et al., 2022), VESPA (Marquet et al., 2022) with pre-computed ProtT5 per-residue embeddings, and ESM-2 (3B) log-odds scores (Lin et al., 2023). For the runtimes of pre-processing steps, see Tab. S4b.

| Preprocessing | | | | |
|---|---|---|---|---|
| **Tool** | **Step** | **Hardware** | **Runtime [s]** | **Memory usage** |
| VespaG | Embedding generation (ESM2) | High-end GPU | 58.6 | 48 GB |
| | | Consumer CPU | 3,226.7 | 20.1 GB |
| | | High-end CPU | 200.73 | 24 GB |
| VESPA | Embedding generation (ProtT5) | High-end GPU | 53.1 | 48 GB |
| GEMME | ColabFold MSA generation | High-end CPU | 870.0 | 17.5 GB |
| | MSA preprocessing | High-end CPU | 37.0 | 17.5 GB |
| | Total | High-end CPU | 907.0 | 17.5 GB |

**Table S5: Runtime of preprocessing steps for methods VespaG, VESPA, and GEMME on 73 unique proteins of the first iteration of the ProteinGym substitution benchmark** (Notin et al., 2023). Hardware used: Consumer CPU: Intel i7-1355U with 12x5 GHz, high-end CPU: 4 cores of a shared AMD EPYC Milan with 32x2.95 GHz, high-end GPU: Nvidia Quattro RTX 8000 or Nvidia RTX 6000 (both 48GB VRAM). The three values reported for VespaG embedding generation correspond to three independent runs on three different kinds of hardware.

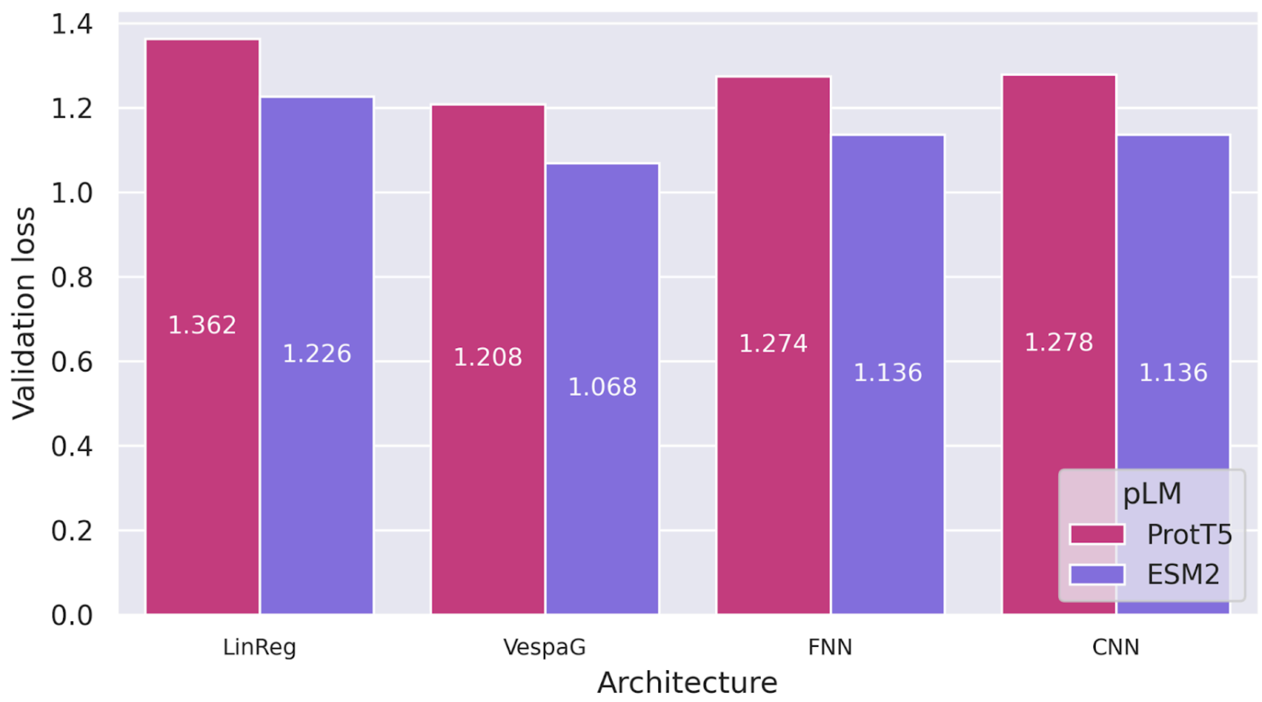

**Figure S1: Influence of the architecture and the input embeddings on VespaG's validation loss.** The loss (mean squared error (MSE) between predicted and target GEMME (Laine et al., 2019) scores) is computed on the randomly selected validation split of the *Hum5k* dataset. ESM-2 embeddings (Lin et al., 2023) yield higher performance than ProtT5 (Elnaggar et al., 2021) across all architectures.

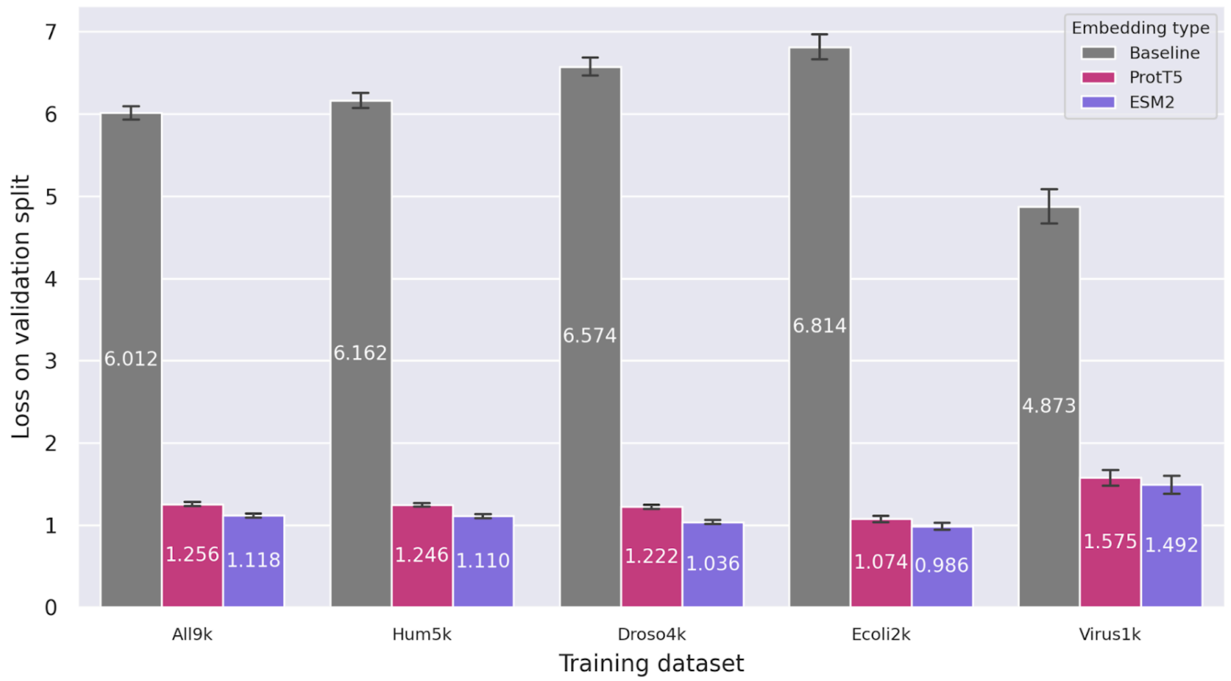

**Figure S2: Influence of the training dataset and the input embeddings on VespaG's validation loss.** The loss (mean squared error (MSE) between predicted and target GEMME (Laine et al., 2019) scores) is computed on the randomly selected validation split of each training dataset and compared to a baseline obtained with randomly permuted GEMME scores (see Materials and Methods). ESM-2 embeddings (Lin et al., 2023) yield significantly higher performance than ProtT5 (Elnaggar et al., 2021) in all cases except the viral training dataset. Both types of embeddings lead to degraded performance on this dataset, whereas the performance of the random baseline is comparatively much better. This observation suggests that GEMME scores have a much lower resolution on viral proteins (more identical or very similar scores). Error bars show 95% confidence intervals.

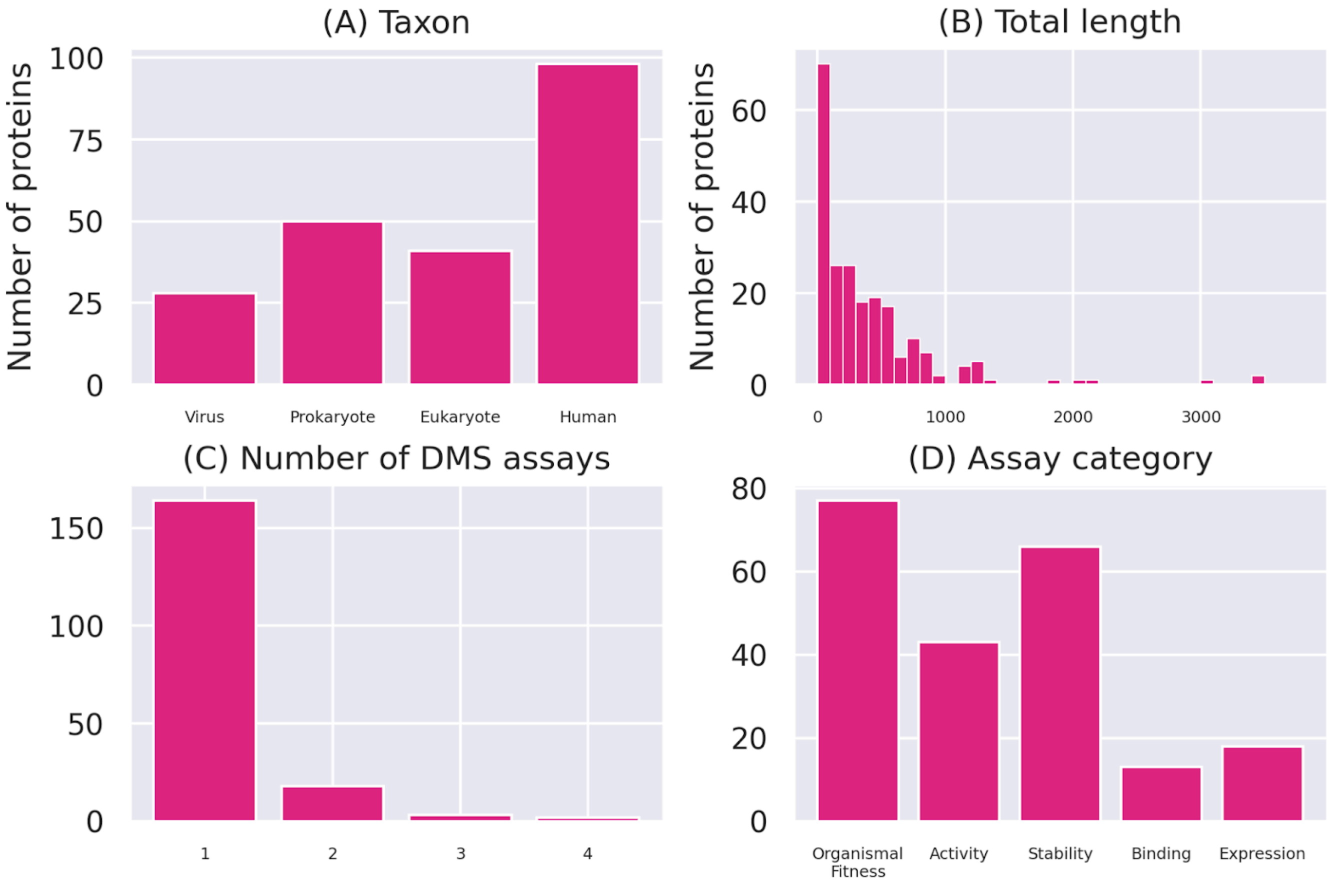

**Figure S3**: **Properties of the ProteinGym substitution benchmark set** (Notin et al., 2023). (A) Distribution of proteins across taxa (Eukaryote referring to non-Human eukaryotic proteins) (B) Distribution of sequence length per protein (C) Distribution of number of deep mutational scanning (DMS) assays per protein (D) Distribution of assay categories.

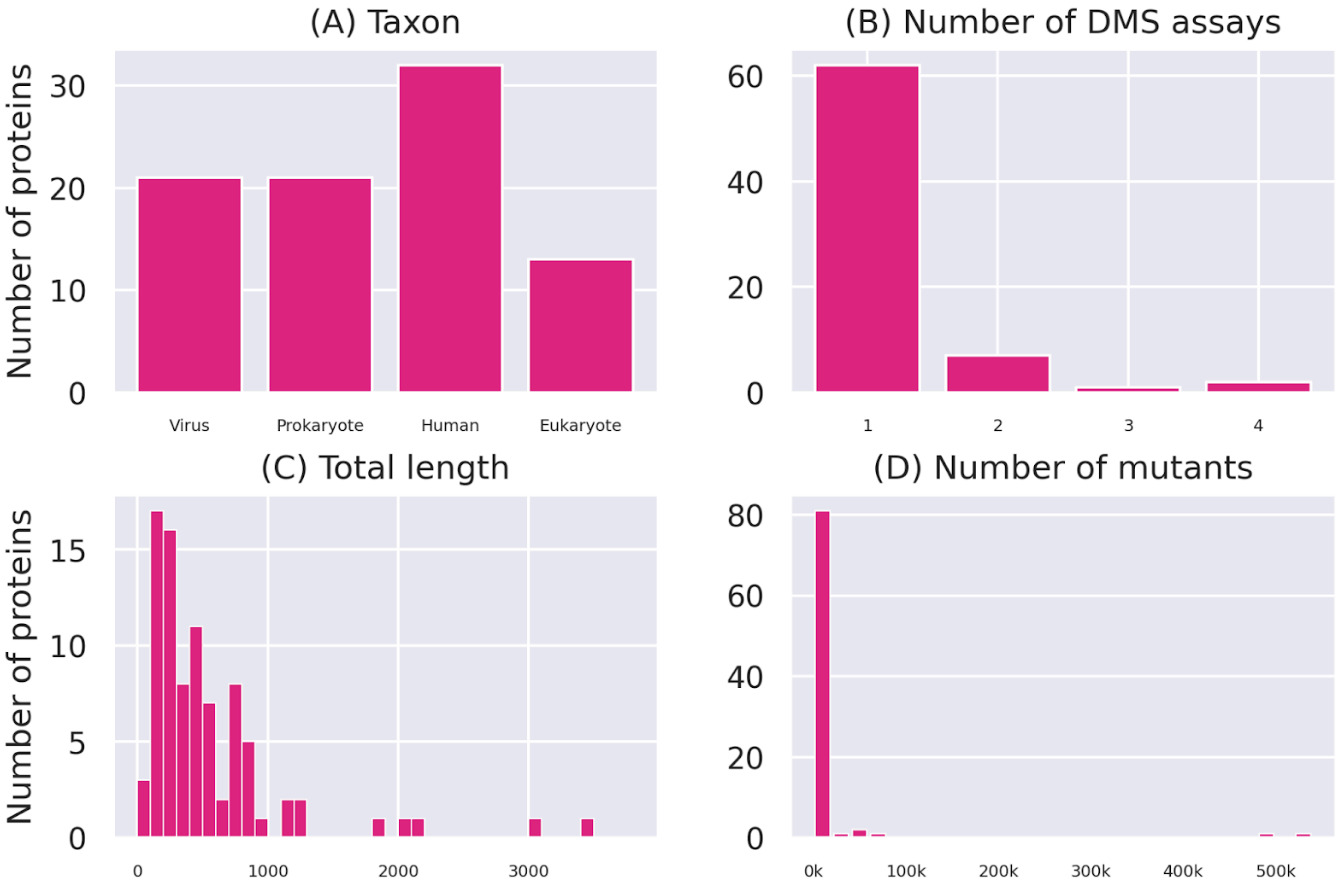

**Figure S4**: **Properties of the first iteration of the ProteinGym substitution benchmark set** (Notin et al., 2023). (A) Distribution of proteins across taxa (Eukaryote referring to non-Human eukaryotic proteins) (B) Distribution of number of deep mutational scanning (DMS) assays per protein (C) Distribution of sequence length per protein (D) Distribution of number of mutants across DMS assays.

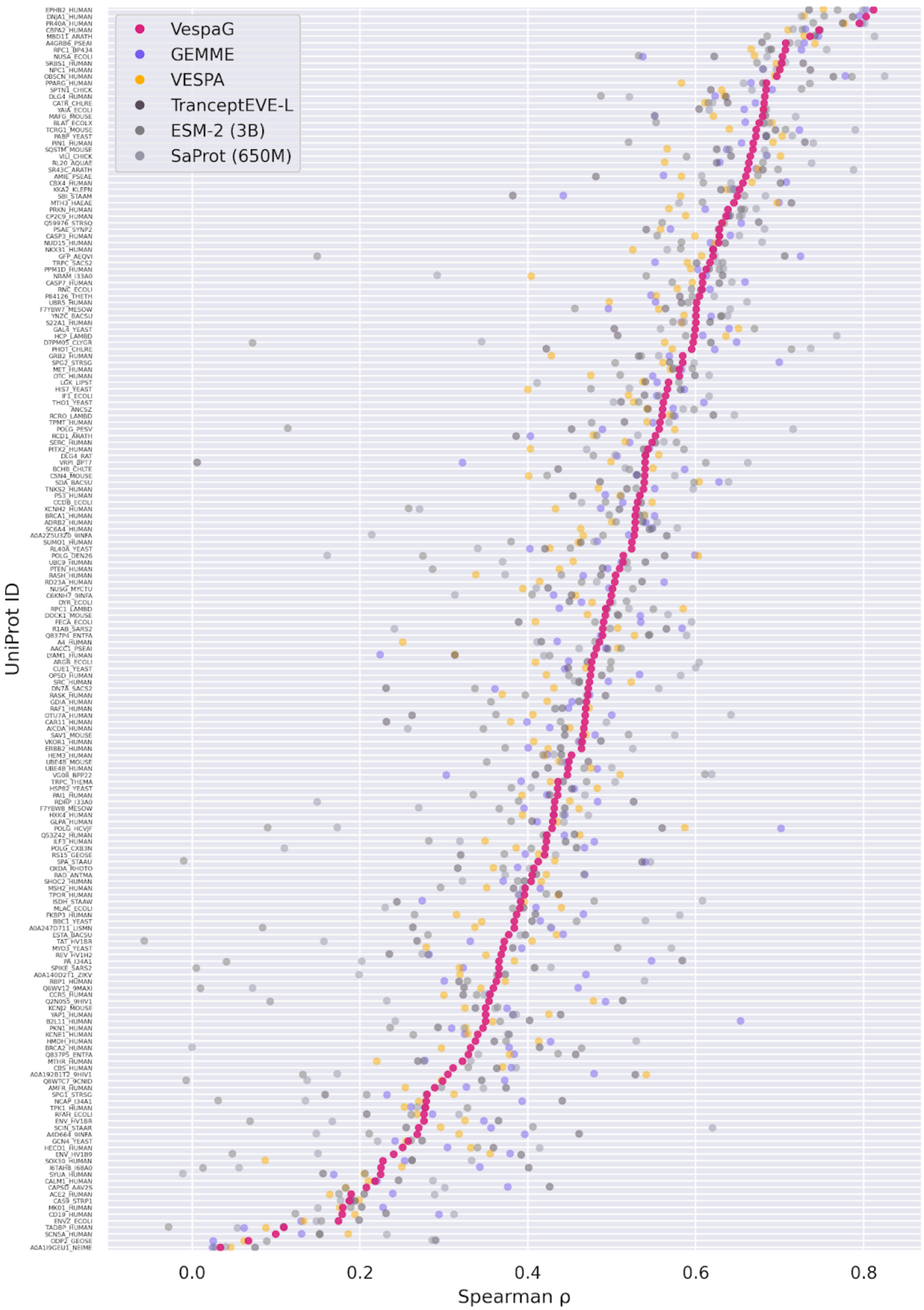

**Figure S5: Average Spearman correlation coefficient ρ between predicted and**

**experimental substitution effect scores for VespaG and SOTA methods on unique proteins of *ProteinGym217*** ordered by VespaG performance. Other methods reported are GEMME (Laine et al., 2019), TranceptEVE L (Notin et al., 2022), VESPA (Marquet et al., 2022), ESM-2 (3B) (Lin et al., 2023), and SaProt (650M) (Su et al., 2024). Spearman correlations of methods depicted in shades of gray (TranceptEVE L, ESM-2 and SaProt) were downloaded from the ProteinGym website.

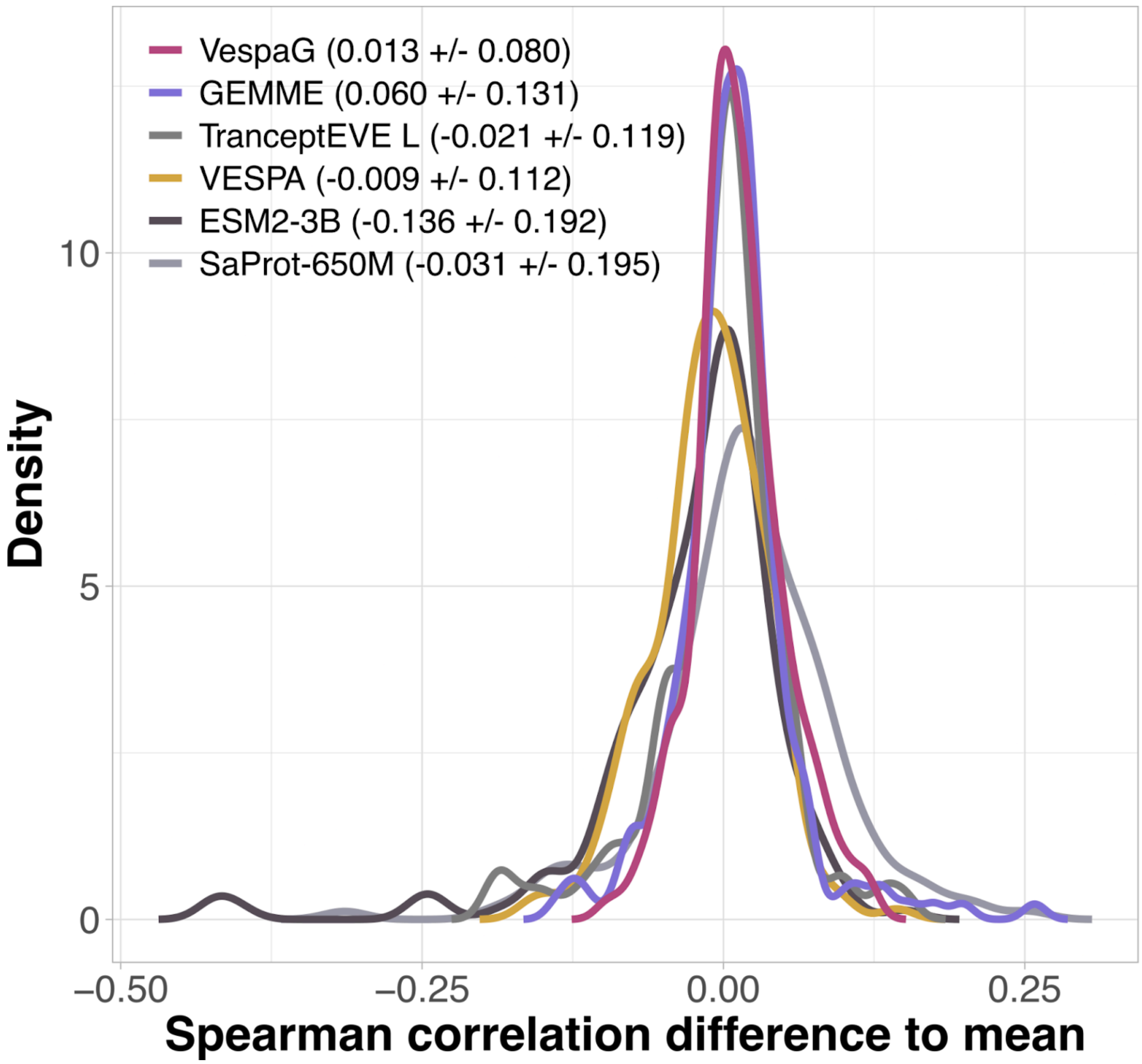

**Figure S6: Consensuality of the predictions on ProteinGymOrganismal189.** For each predictor, we report the distribution of the differences between its performance values (Spearman correlation coefficients with experiments) and the average performance of all six highlighted predictors on the *ProteinGymOrganismal189* test set. The mean and standard deviation for each density are indicated in parenthesis.

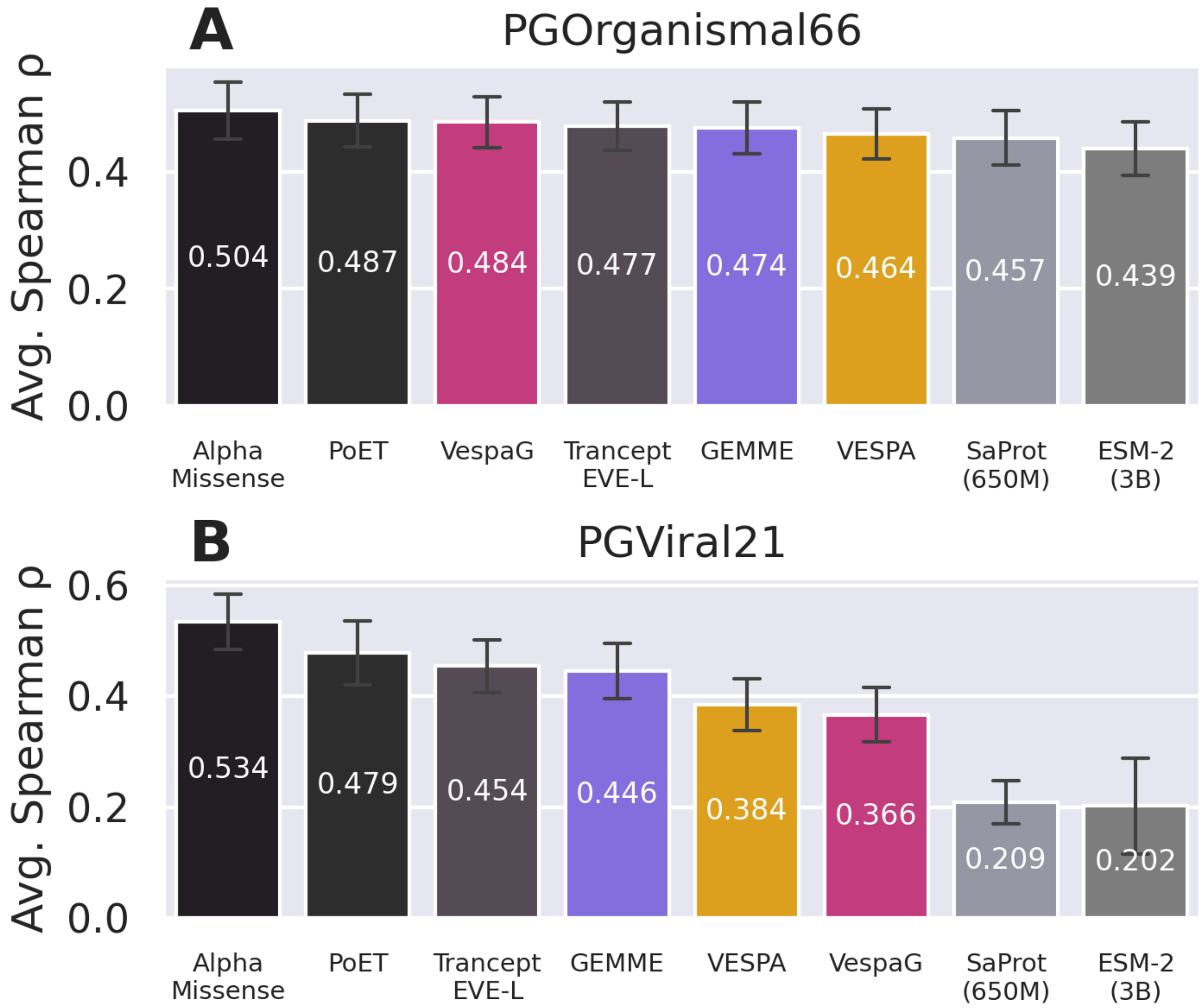

**Figure S7: Average Spearman correlation coefficient ρ between predicted and experimental substitution effect scores of VespaG and SOTA methods on the first iteration of ProteinGym.** Methods presented are VespaG, GEMME (Laine et al., 2019), TranceptEVE L (Notin et al., 2022), VESPA (Marquet et al., 2022), ESM-2 (3B) (Lin et al., 2023), SaProt (650M) (Su et al., 2024), AlphaMissense (Cheng et al., 2023), and PoET (Truong Jr and Bepler, 2023). Results for methods depicted in gray were downloaded from ProteinGym (TranceptEVE L, ESM-2 and SaProt) or taken from the respective publications (AlphaMissense, PoET). Panel (A) *ProteinGymOrganismal66*, containing 66 experimental assays for 54 eukaryotic and prokaryotic proteins; (B) *ProteinGymViral21*, containing 21 assays for 19 viral proteins of the first iteration of the ProteinGym substitution benchmark.

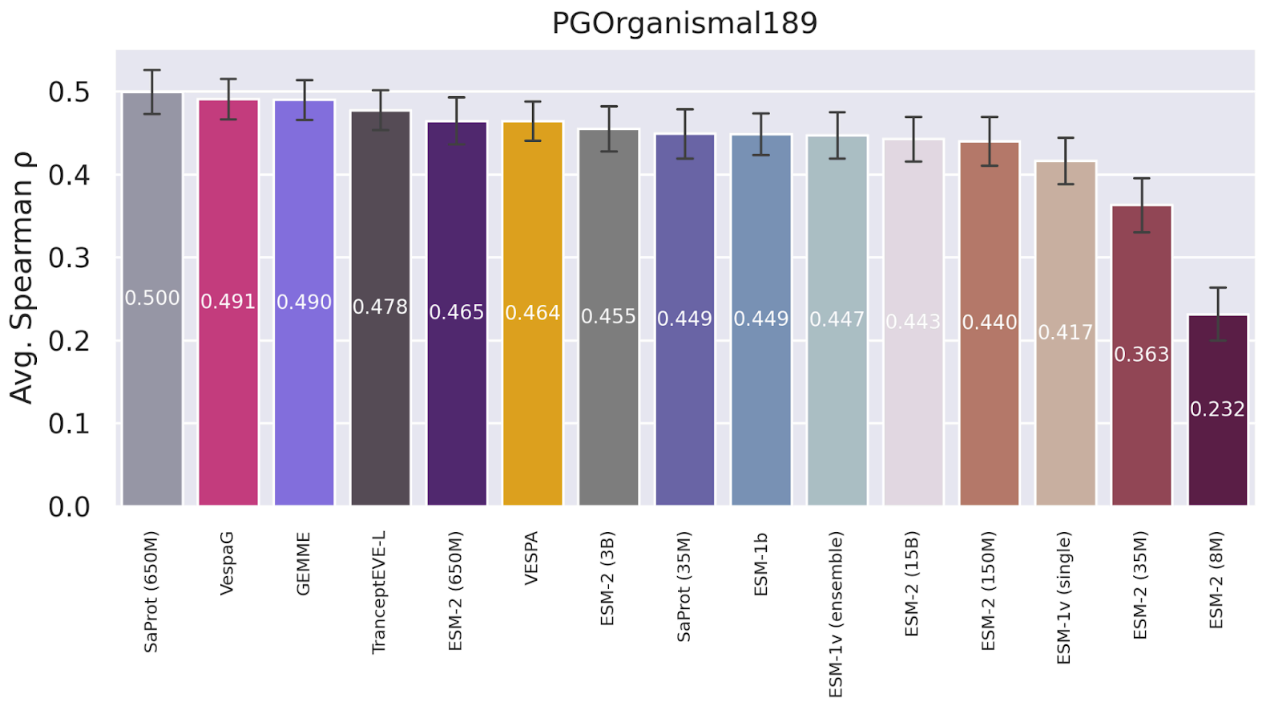

**Figure S8: Average Spearman correlation coefficient ρ between experimental and predicted substitution effect scores of VespaG and SOTA methods on 189 organismal DMS assays of the ProteinGym substitution benchmark.** Methods presented are VespaG, GEMME (Laine et al., 2019), TranceptEVE L (Notin et al., 2022), VESPA (Marquet et al., 2022), 6 variants of ESM-2 (Lin et al., 2023), 2 variants of SaProt (Su et al., 2024), 2 variants of ESM-1v (Meier et al., 2021), and ESM-1b (Rives et al., 2021). Results for all methods except VespaG, GEMME, and VESPA were downloaded from ProteinGym.

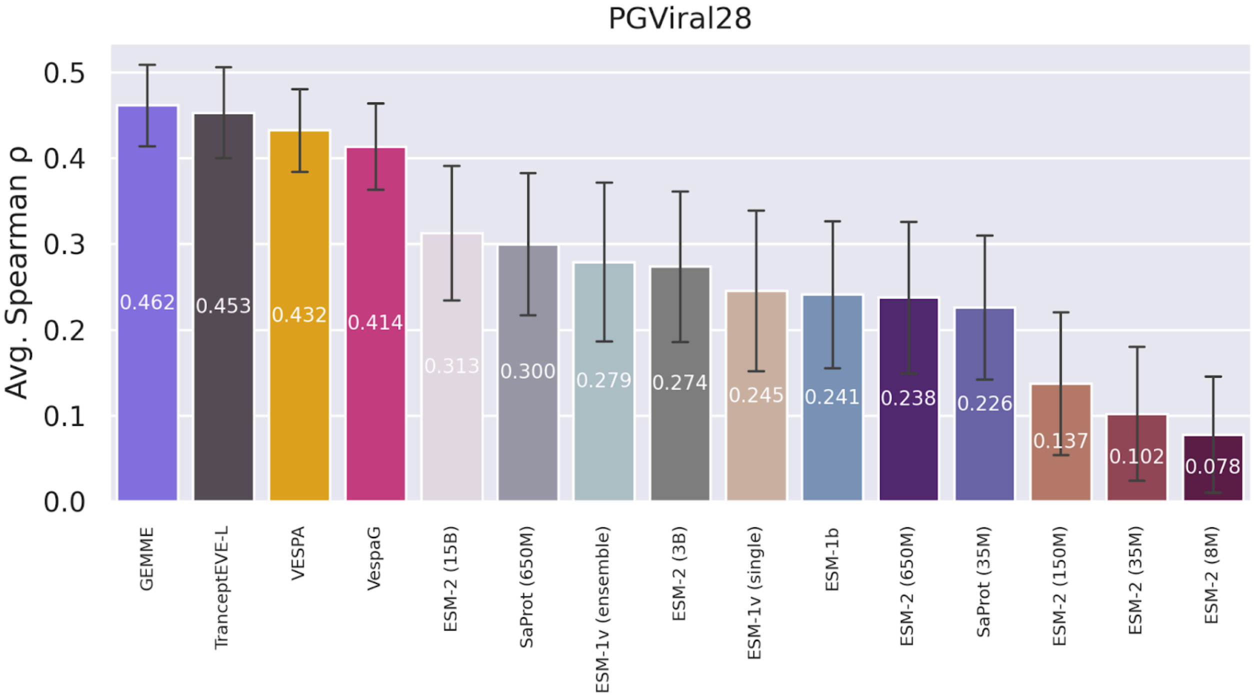

**Figure S9: Average Spearman correlation coefficient ρ between experimental and predicted substitution effect scores of VespaG and SOTA methods on 28 viral DMS assays of the ProteinGym substitution benchmark.** Methods presented are VespaG, GEMME (Laine et al., 2019), TranceptEVE L (Notin et al., 2022), VESPA (Marquet et al., 2022), 6 variants of ESM-2 (Lin et al., 2023), 2 variants of ESM-1v (Meier et al., 2021), and ESM-1b (Rives et al., 2021). Results for all methods except VespaG, GEMME, and VESPA were downloaded from ProteinGym.

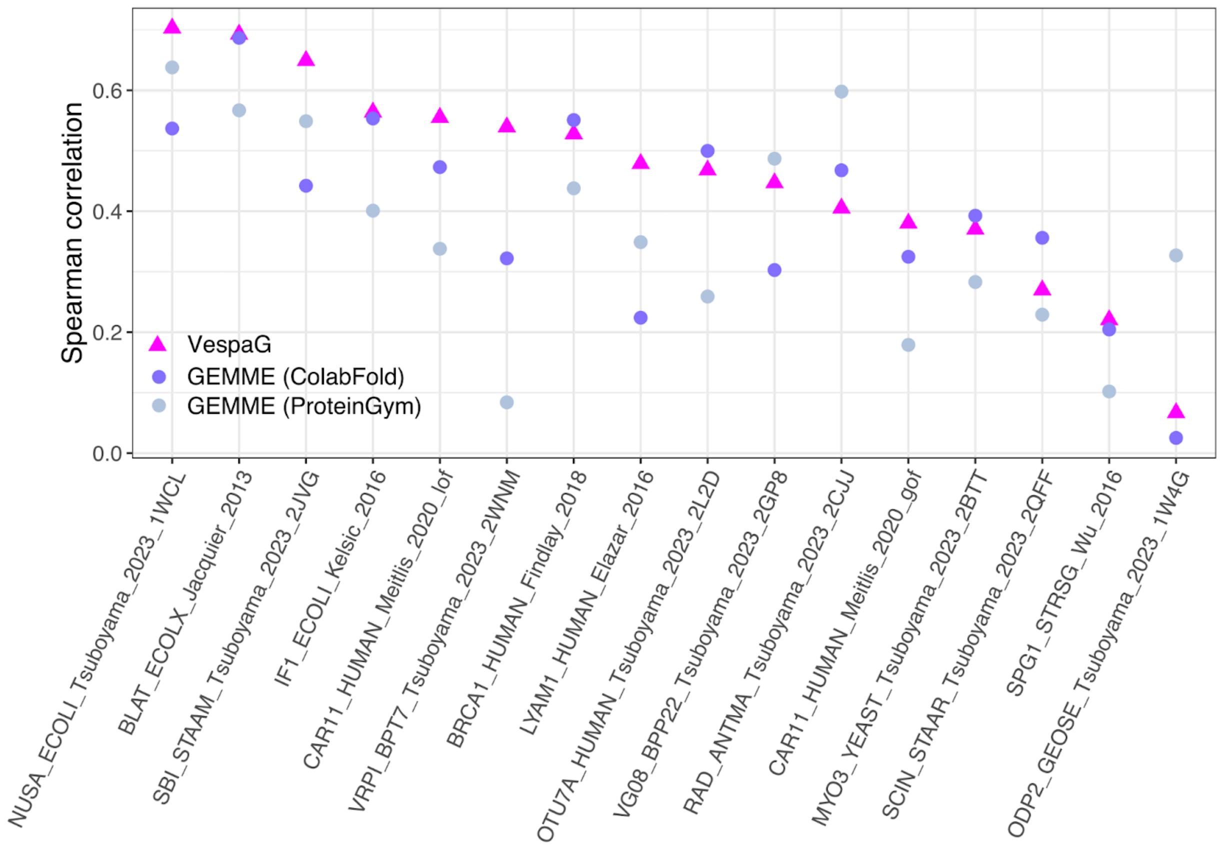

**Figure S10: Comparison of Spearman correlation depending on the input alignment.** We consider the subset of DMS where GEMME Spearman correlation varied by more than 0.1 between two different input alignment generation protocols, namely the MMseqs2-based strategy implemented in ColabFold (this work) (Laine et al., 2019; Mirdita et al., 2022) and the JackHMMER-based strategy (Johnson et al., 2010) implemented in ProteinGym.

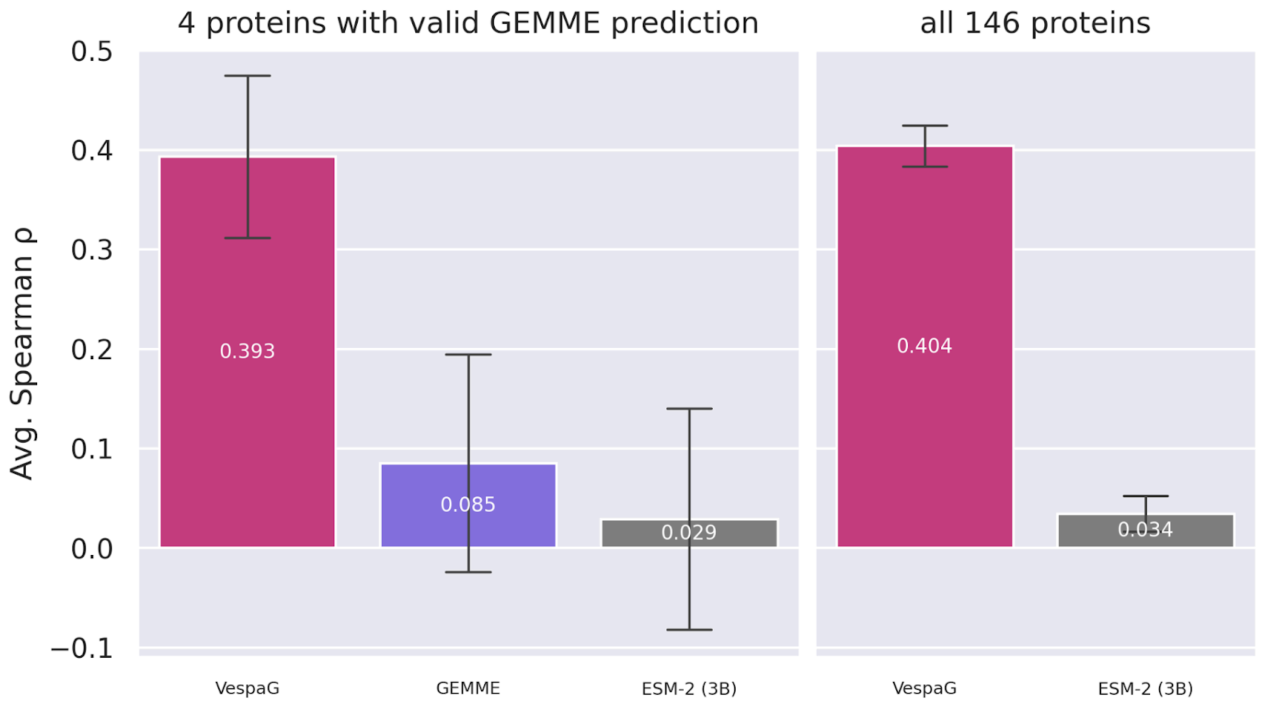

**Figure S11: Average Spearman correlation coefficient ρ between predicted mutational effect scores and experimental ΔΔG scores of VespaG, GEMME (Laine et al., 2019), and ESM-2 (3B) (Lin et al., 2023) on the StabilityDeNovo146 dataset (Tsuboyama et al., 2023).** The left subplot shows four proteins for which GEMME could produce predictions, and the right subplot excluding GEMME shows all 146 proteins.
